# Opponent learning with different representations in the cortico-basal ganglia circuits

**DOI:** 10.1101/2021.10.29.466375

**Authors:** Kenji Morita, Kanji Shimomura, Yasuo Kawaguchi

## Abstract

The direct and indirect pathways of the basal ganglia (BG) have been suggested to learn mainly from positive and negative feedbacks, respectively. Since these pathways unevenly receive inputs from different cortical neuron types and/or regions, they may preferentially use different state/action representations. We explored whether such combined use of different representations coupled with appetitive or aversive learning has computational benefits. We simulated reward learning tasks in dynamic environments, and examined the performance of animal modeled as an agent equipped with two learning systems, each of which used individual representation (IR) or successor representation (SR) of states. With varying the combination of IR or SR and also the learning rates from positive and negative reward prediction errors (RPEs) in each system, we found that combination of an SR-based system learning mainly from positive RPEs and an IR-based system learning mainly from negative RPEs could achieve good performance, as compared to other combinations, in many situations. The architecture of such a combination provides a novel coherent explanation for the functional significance and underlying mechanism of diverse findings about the cortico-BG circuits. These results suggest that combining different representations with appetitive and aversive learning is an effective learning strategy adopted by the brain.

## Introduction

In the standard reinforcement learning (RL), updates based on positive reward prediction errors (RPEs) and those based on negative RPEs are executed in the same manner. However, in the brain, there appear to exist distinct neural circuits that are specialized for appetitive or aversive learning. Specifically, a number of findings have suggested or implied that the direct and indirect pathways of the basal ganglia (BG), originating from the striatal projection neurons (SPNs) expressing D1- and D2-type dopamine (DA) receptors (D1Rs and D2Rs), are potentiated by positive and negative feedbacks, respectively (Frank et al., 2004; Hikida et al., 2010; Iino et al., 2020; Kim et al., 2017; Kravitz et al., 2012; Lee et al., 2021; Tai et al., 2012). There are also studies suggesting that distinct circuits involving the orbitofrontal cortex (OFC) operate for appetitive and aversive feedback-based learning (Groman et al., 2019a). Computational works suggest that dual learning systems can realize estimation of costs and benefits (Collins and Frank, 2014; Möller and Bogacz, 2019), as well as estimation of not only the mean but also the uncertainty of rewards (Mikhael and Bogacz, 2016).

These existing dual learning-system models assume that both systems use the same way of representation of states or actions. Theoretically, various ways of representation can be considered, and different brain regions or neural populations may generally use different representations (Chen et al., 2014; Town et al., 2017; Wang et al., 2020). While there is evidence that the two BG pathways receive inputs from same types of corticostriatal neurons (Ballion et al., 2008; Kress et al., 2013), it has also been suggested that different neuron types (Lei et al., 2004; Morita, 2014; Reiner et al., 2010) and/or cortical areas (Lu et al., 2021; Wall et al., 2013) may not evenly target/activate these pathways. As for the suggested distinct OFC circuits for appetitive and aversive learning, the suggested circuits are the amygdala-OFC and OFC-nucleus accumbens (NAc) pathways, respectively (Groman *et al*., 2019a). Therefore, in both cases, it is conceivable that, between the appetitive and aversive learning systems, states/actions are represented by at least partially different neural regions or populations, and thus in different styles.

There are two largely different ways of state/action representation (Sutton and Barto, 1998). One is to represent each individual state/action separately. This simplest representation (or equivalent ones) has been explicitly or implicitly assumed to exist in the brain RL circuits in many previous studies. The other is to represent each state/action by a set of features (e.g., represent a point in a two-dimensional space by a set of *x*- and *y*-coordinates). Among various ways of feature-based representations, recent studies (Garvert et al., 2017; Momennejad et al., 2017; Russek et al., 2017; Russek et al., 2021; Stachenfeld et al., 2017) suggest that representation of states by their successors, named the successor representation (SR) (Dayan, 1993), may be used in the brain. SR contains information about state transitions in the environment under a given policy, and thereby enables the agent to quickly adapt to changes in distant outcomes through temporal difference (TD) RPE-based learning, without using an explicit model of the environment. It can thus beautifully explain (Russek *et al*., 2017) the empirical suggestions that both (apparently) model-based behavior and model-free or habitual behavior are controlled by the DA-cortico-BG systems although different portions appear to be responsible (Balleine and O’Doherty, 2010; Dolan and Dayan, 2013) by assuming that SR and individual (punctate) representation (IR) are respectively used.

Given these suggestions and considerations, it seems possible that there exist two neural systems, which may differently learn from positive and negative RPEs and may adopt different ways of state representations. In the present study, we examined its possible consequences through simulations. Specifically, we modeled animal as an agent equipped with two learning systems, and simulated its behavior in reward learning tasks in dynamic environments, with varying the adopted representation, SR or IR, and the ratio of learning rates for positive versus negative TD-RPEs in each system.

## Results

### Performance of combined SR-based and IR-based systems, with learning rate ratio varied

We first simulated a reward navigation task in a two-dimensional grid space, in which reward location changed over time (Figure 1), and examined the performance of an agent consisting of two systems, which adopted SR or IR and may have different ratios of learning rates for positive versus negative TD-RPEs; the two systems developed system-specific state values, and their average (named the integrated state value) was used for action selection and TD-RPE calculation (Figure 2) (see the Materials and Methods for details). We first examined the case in which one system employed the SR whereas the other adopted the IR, systematically varying the ratio of learning rates from positive and negative TD-RPEs for each system, denoted as *α*_SR+_ / *α*_SR−_ for the SR-based system and *α*_IR+_ / *α*_IR−_ for the IR-based system. The sums of the learning rates from positive and negative TD-RPEs (i.e., *α*_SR+_ + *α*_SR−_ and *α*IR+ + *α*_IR−_) were kept constant at 1, and the inverse temperature and the time discount factor were also kept constant at *β* = 5 and *γ* = 0.7. We counted how many rewards the agent obtained during the reward navigation task as a measure of performance, and examined how the obtained rewards, averaged across 100 simulations for each condition, varied depending on the learning rate ratios in both systems.

**Figure 1.**
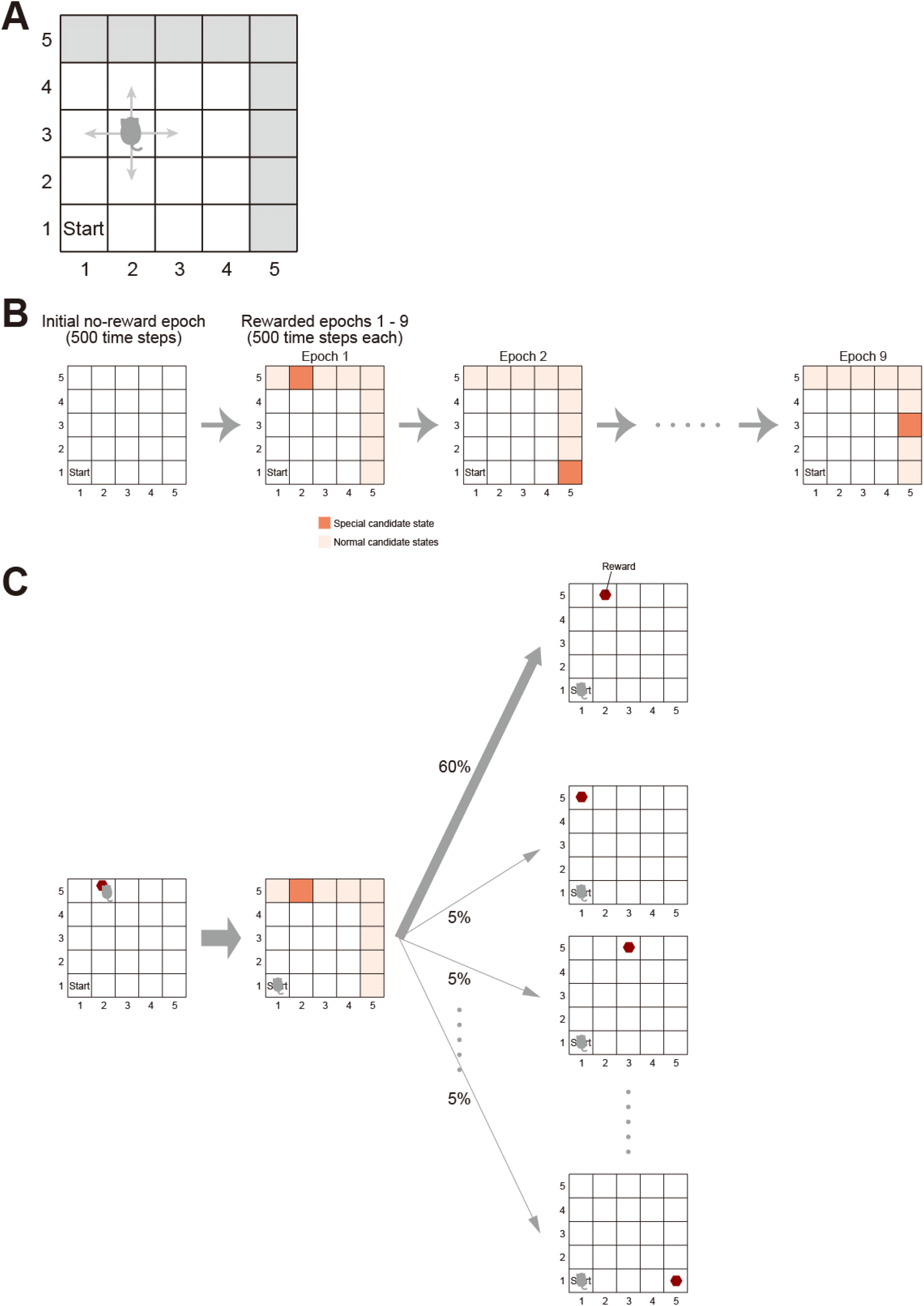
The simulated reward navigation task. **(A)** The 5 × 5 grid space, where the agent moved around. The agent started from the fixed start state, (1, 1), and moved to one of the neighboring states (two states at the corners, three states at the edges, and four states elsewhere) at each time step. There were nine reward candidate states, where reward could potentially be placed, namely, (1, 5), (2, 5), (3, 5), (4, 5), (5, 1), (5, 2), (5, 3), (5, 4), and (5, 5) (indicated by the gray color). **(B)** Epochs in the task. During the initial 500 time steps, there was no reward, and this was called the no-reward epoch. During the next 500 time steps, one of the nine reward candidate states was specified as a special candidate state, whereas the remaining eight reward candidate states were regarded as normal candidate states. There were in total nine rewarded epochs (500 time steps for each), and each one of the nine reward candidate states became the special candidate state in one of the nine epochs; the order was determined by pseudorandom permutation in each single simulation. **(C)** In the rewarded epochs, if the agent reached the rewarded state and obtained the reward, the agent was immediately carried back to the start state, and a new reward was introduced into a state, which was the special reward candidate state with 60% probability and one of the eight normal candidate states with 5% probability for each.

**Figure 2.**
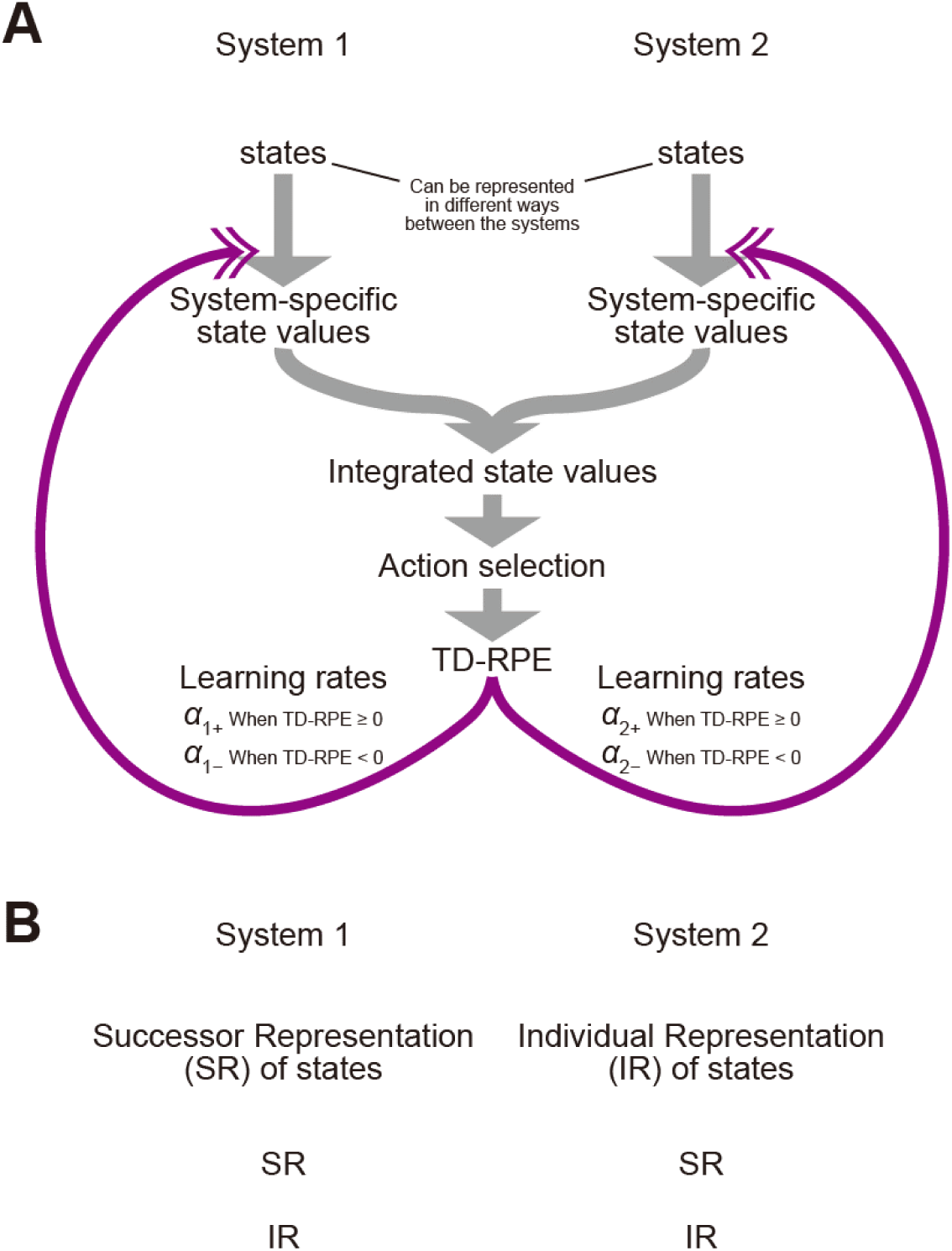
The reinforcement learning model with two systems. **(A)** Model architecture. The model consists of two learning systems, system 1 and system 2, which may use different ways of state representations. Each system has its own system-specific value of each state, and their mean (average) becomes the integrated state value. The agent selects an action to move to a neighboring state depending on their integrated values in a soft-max manner. The integrated values are also used for calculating the temporal-difference reward prediction error (TD-RPE). The TD-RPE was then used for updating the system-specific state values in each of the two systems. The leaning rates of the updates of the system-specific state values can differ between the two systems and also depending on whether the TD-RPE is positive (non-negative) or negative. **(B)** Combinations of the ways of state representation in the two systems. Combination of the successor representation (SR) in system 1 and the individual representation (IR) in system 2 was examined in Figures 3-5, 7, 8, S1, S3, and Table S1. Cases with only the SR in both systems were examined in Figures 6, 8, S2, Tables S1 and S2. Cases with only the IR in both systems were examined in Figures 6, 8, S2, S3, and Table S1.

Figure 3A shows the results. As shown in the figure, the performance greatly varied depending on the conditions, and the best performance was achieved in the conditions in which *α*_SR+_ / *α*_SR−_ was high (larger than 1) and *α*_IR+_ / *α*_IR−_ was low (smaller than 1), i.e., the SR-based system learned mainly from positive TD-RPEs whereas the IR-based system learned mainly from negative TD-RPEs. Figure 3B shows the standard deviation (SD) and standard error of the mean (SEM) for the conditions where *α*_SR+_ / *α*_SR−_ times *α*_IR+_ / *α*_IR−_ was equal to 1 (i.e., the conditions on the horizontal diagonal in Figure 3A), and Figure 3C shows the frequency (number of times) that each combination of *α*_SR+_ / *α*_SR−_ and *α*_IR+_ / *α*_IR−_ gave the best mean performance over 100 simulations when 100 simulations for each combination were executed 50 times (including the one shown in Figure 3A,B). As shown in these figures, the peak of the performance was rather broad, but was mostly in the range where *α*_SR+_ / *α*_SR−_ was large and *α*_IR+_ / *α*_IR−_ was small. We also examined the cases where the sums of the learning rates from positive and negative TD-RPEs were increased (1.25) or decreased (0.75), and also the inverse temperature and the time discount factor were varied (*β* = 5 or 10 and *γ* = 0.7 or 0.8). As shown in Figure 4, the result that combination of the SR-based system learning mainly from positive TD-RPEs and the IR-based system learning mainly from negative TD-RPEs achieved the best performance was preserved across these parameter changes.

**Figure 3.**
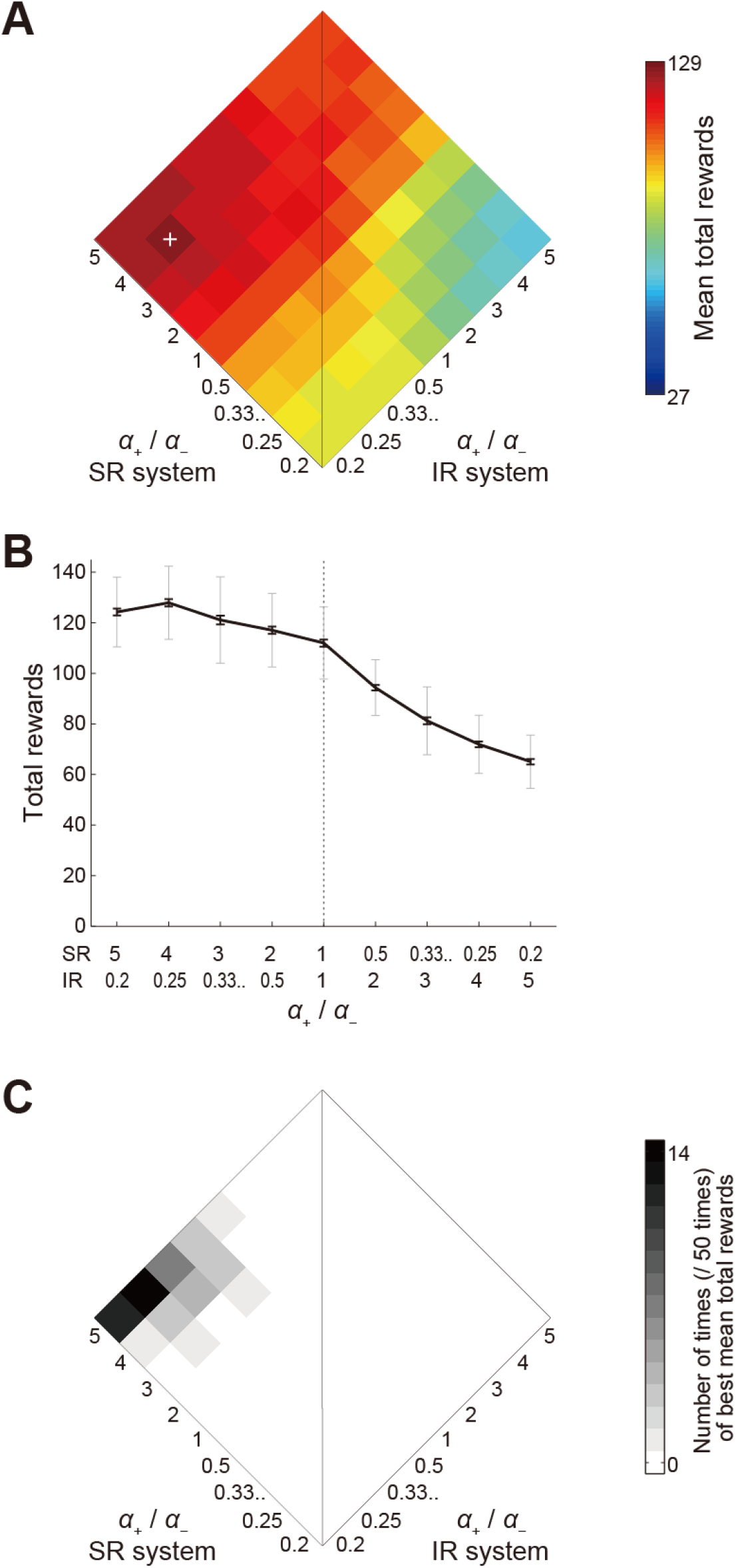
Performance of the model consisting of a system using the SR and another system using the IR, and its dependence on the ratios of the learning rates from positive and negative TD-RPEs in each system. **(A)** Mean performance over *n* = 100 simulations for each condition. The axis rising to the left indicates the ratio of positive- / negative-error-based learning rates (denoted as *α*_+_/*α*_−_) in the system using the SR, while the axis rising to the right indicates the same ratio in the system using the IR. The sum of the learning rates from positive and negative TD-RPEs (*α*_+_ + *α*_−_) in each system was constant at 1 in any conditions shown in this figure. The inverse temperature *β* was 5, and the time discount factor *γ* was 0.7. The vertical line corresponds to conditions where the *α*_+_/*α*_−_ ratio is equal for both systems (bottom: negative error-based learning dominates; top: positive error-based learning dominates). The left side of the vertical line corresponds to conditions where the *α*_+_/*α*_−_ ratio is larger in the SR-based system than in the IR-based system, whereas the opposite is the case for the right side of the vertical line. The color in each square pixel indicates the mean total obtained rewards in the task, averaged across 100 simulations, for each condition (i.e., set of *α*_SR+_/*α*_SR−_ and *α*_IR+_/*α*_IR−_ at the center of the square pixel), in reference to the rightmost color bar. The white cross indicates the set of *α*_+_/*α*_−_ ratios that gave the best performance (*α*_+_/*α*_−_ = 4 and 0.25 for the SR- and IR-based systems, respectively) among those examined in this figure. Note that the minimum value of the color bar is not 0. Also, the maximum and minimum values of the color bar do not match the highest and lowest performances in this figure; rather, they were set so that the highest and lowest performances in all the simulations of the same task with different parameters and/or model architecture (i.e., not only SR+IR but also SR+SR and IR+IR) shown in this figure and Figures 4 and 6 can be covered. **(B)** The black solid line, gray thin error-bars, and black thick error-bars respectively show the mean, standard deviation (SD, normalized by *n*; same hereafter), and standard error of the mean (SEM, approximated by SD/√*n*; same hereafter) of the performance over *n* = 100 simulations for the conditions where *α*_+_/*α*_−_ for SR system times *α*_+_/*α*_−_ for IR system was equal to 1 (i.e., the conditions on the horizontal diagonal in (A)). **(C)** Frequency (number of times) that each condition gave the best mean performance over 100 simulations when 100 simulations for each condition were executed 50 times (including the one shown in (A,B)).

**Figure 4.**
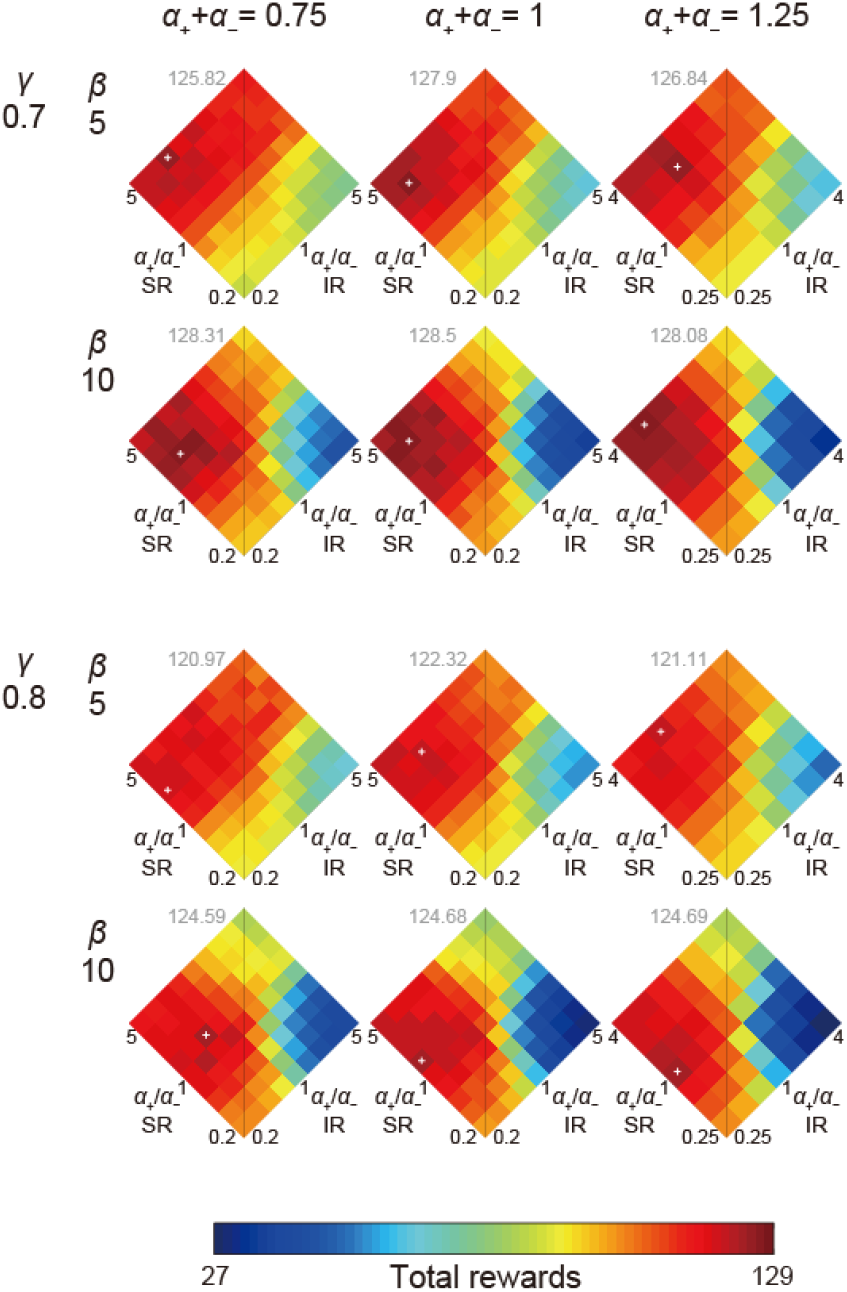
Performance of the model consisting of an SR-based system and an IR-based system with various sets of parameters. The sum of the learning rates from positive and negative TD-RPEs (*α*_+_ + *α*_−_) was varied over 0.75, 1, and 1.25. The inverse temperature *β* was varied over 5 and 10. The time discount factor *γ* was varied over 0.7 and 0.8. The ratio of positive- / negative-error-based learning rates (*α*_+_/*α*_−_) was varied over 0.2, 0.25, 1/3, 0.5, 1, 2, 3, 4, and 5 for the cases with *α*_+_ + *α*_−_ = 0.75 or 1, but 0.2 and 5 were omitted for the cases with *α*_+_ + *α*_−_ = 1.25 in order to avoid learning rate larger than 1. The color-performance correspondence is the same as in Figure 3. The white cross in each panel indicates the set of *α*_+_/*α*_−_ ratios that gave the best performance among those examined in that condition, and the gray number near the top of each panel indicates the best performance (total rewards). The panel with *α*_+_ + *α*_−_ = 1, *β* = 5, and *γ* = 0.7 shows the same results as shown in Figure 3.

### Learning profiles of the model with different combinations of the learning rate ratios

Going back to the case with original parameter values used in Figure 3 (*α*_+_ + *α*_−_ = 1, *β* = 5, and *γ* = 0.7), we examined the learning curves of the model in the case where the ratio of positive versus negative TD-RPE-based learning rate was respectively high (4) and low (0.25) in the SR-based and IR-based systems (i.e., the case achieving good performance), compared with the cases where the ratio was 1 in both systems or the ratio was low (0.25) and high (4) in the SR-based and IR-based systems. Figure 5A shows the mean learning curves in each case, i.e., the mean time (number of time steps) used for obtaining the first, second, third,.. reward *placed* in each rewarded epoch, averaged across eight (from the second to the ninth) rewarded epochs and also across simulations (if a reward was placed in the first epoch and obtained in the second epoch, for example, it was regarded as the last reward in the first epoch rather than the first reward in the second epoch). Only the cases in which reward was obtained in not smaller than a quarter of (25 out of total 100) simulations were plotted, in order to see the general average behavior of the model. Comparing the case with high *α*_SR+_ / *α*_SR− &_ low *α*_IR+_ / *α*_IR−_ (brown line) and the case with low *α*_SR+_ / *α*_SR− &_ high *α*_IR+_ / *α*_IR−_ (blue-green line), the time consumed for obtaining the first reward in an epoch, as well as the asymptotic value of the time spent for reward acquisition, were much longer in the latter case. The case with equal ratios for both systems (red line) showed an intermediate behavior.

**Figure 5.**
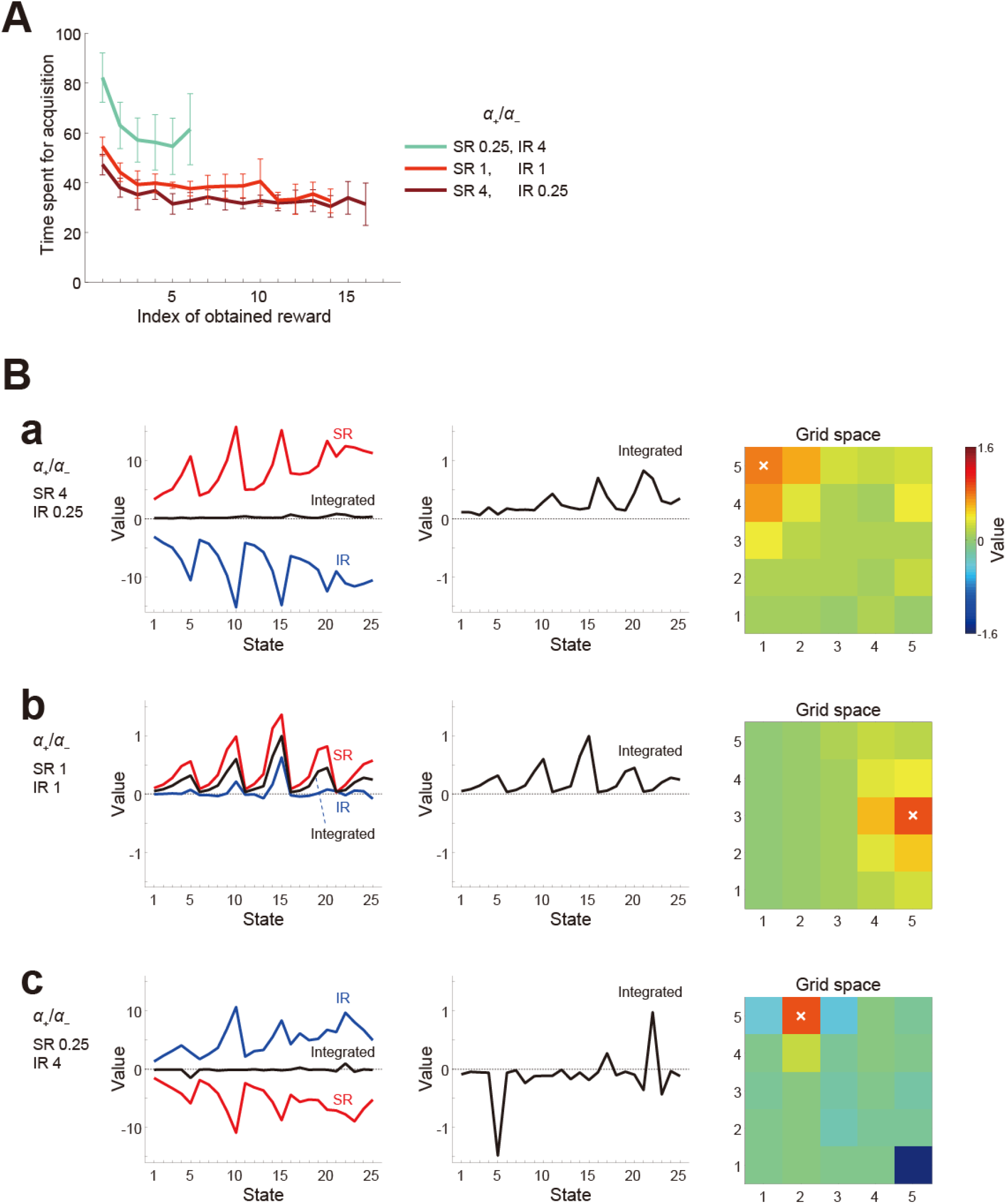
Learning profiles of the model consisting of an SR-based system and an IR-based system with different combinations of the ratios of positive- / negative-error-based learning rates in each system. Three cases included in Figure 3 (where the sum of the learning rates from positive and negative TD-RPEs (*α*_+_ + *α*_−_) in each system was 1, the inverse temperature *β* was 5, and the time discount factor *γ* was 0.7) were analyzed. **(A)** Mean learning curves. The curves indicate the mean time (number of time steps) used for obtaining the first, second, third,.. reward placed in each rewarded epoch (horizontal axis), averaged across simulations and also across eight (from the second to the ninth) rewarded epochs. Only the cases in which reward was obtained in not smaller than a quarter of (25 out of total 100) simulations in all of the eight epochs were plotted. The brown, red, and blue-green curves correspond to the conditions with (*α*_+_/*α*_−_ for SR-system, *α*_+_/*α*_−_ for IR-system) = (4, 0.25), (1, 1), and (0.25, 4), respectively (the colors match those in the corresponding pixels in Figure 3). The error-bars indicate ±SD across the eight epochs (after taking the averages across simulations). **(B)** Examples of the system-specific state values and the integrated state values. Panels in (a), (b), and (c) correspond to the conditions with (*α*_+_/*α*_−_ for SR-system, *α*_+_/*α*_−_ for IR-system) = (4, 0.25), (1, 1), and (0.25, 4), respectively. The left panels show the system-specific state values (red line: SR-based system, blue line: IR-based system) and the integrated state values (black solid line). The horizontal axis indicates 25 states (1-5 correspond to (1,1)-(5,1) in Figure 1/5B-right; 6-10 correspond to (1,2)-(5,2), and so on). Values just after the agent obtained the last reward in the last (9th) rewarded epoch (indicated by the white crosses in the right panels) in single simulations, in which that reward was placed at the “special candidate state (with high probability of becoming a rewarded state: see the Materials and Methods)” in that epoch, were shown. Note the differences in the vertical scales between (a) or (c) and (b). The middle panels show only the integrated state values. The vertical axes in (a) and (c) were enlarged as compared to the left panels. The right panels also show the integrated state values as a heat map on the 5 × 5 grid space.

We also looked at how each system developed system-specific values of the states, again in the three cases where *α*_+_ / *α*_−_ for the SR-based and IR-based systems were 4 and 0.25, 1 and 1, or 0.25 and 4. Specifically, we looked at single-simulation examples of those values just after the agent obtained the last reward in the last (9th) rewarded epoch in single simulations, in which that reward was placed at the “special candidate state (with high probability of becoming a rewarded state: see the Materials and Methods)” in that epoch, in the three cases. Figure 5Ba shows the good-performance case where *α*_SR+_ / *α*_SR−_ was high (4) and *α*_IR+_ / *α*_IR−_ was low (0.25). The left panels show both the system-specific and integrated values, and the middle and right panels show only the integrated values in different ways. As shown in the figure, the system-specific state values of the SR-based system (red line in Figure 5Ba left) were all positive whereas those of the IR-based system (blue line) were all negative, and these two look largely symmetric. This seems in line with the observation (Cui et al., 2013) that the striatal direct- and indirect-pathway neurons showed concurrent activation. Meanwhile, the average of these two system-specific values, i.e., the integrated state values (black lines in Figure 5Ba left and middle; also shown as a heat map in Figure 5Ba right), have much smaller magnitudes but show a clear pattern, which is expected to approximate the true state values under the policy that the agent was taking. Figure 5Bb shows the case where both *α*_SR+_ / *α*_SR−_ and *α*_IR+_ / *α*_IR−_ were 1. In this case, both systems developed similar values. Figure 5Bc shows the case where *α*_SR+_ / *α*_SR−_ was low (0.25) and *α*_IR+_ / *α*_IR−_ was high (4). In this case, the SR- and IR-system-specific values were negative and positive, respectively, in contrast to the case of Figure 5Ba with high *α*_SR+_ / *α*_SR−_ and low *α*_IR+_ / *α*_IR−_. The integrated values have comparable magnitudes to the other two cases, but were negative in many states. Comparing the right panels of Figure 5Ba and Figure 5Bc, the peak of the value function looks sharper in the latter, presumably reflecting the smaller degree of positive TD-RPE-dependent learning in the SR-based system.

### Cases where both systems employ only the IR or only the SR

We also examined the cases where both systems employed only the SR, or only the IR. Figure 6A and 6B show the performance results for the SR only and IR only cases, respectively, for the same sets of parameters as shown in Figure 4 for the case of SR and IR combination. Notably, if the ratio of positive versus negative TD-RPE-based learning rates was equal between both systems that employ only the SR or IR, the two systems behaved in exactly the same manner, and thus such conditions (on the vertical lines in Figure 6A and 6B) were equivalent to having only a single system. Also, Figure 6A and 6B only show either the left or right side, because “*α*_1+_ / *α*_1−_ = 0.2 and *α*_2+_ / *α*_2−_ = 3”, for example, are equivalent to “*α*_1+_ / *α*_1−_ = 3 and *α*_2+_ / *α*_2−_ = 0.2” given that both systems 1 and 2 employed the same representation (SR or IR) so that we examined only one of these, and Figure 6A shows the left side whereas Figure 6B shows the right side just because such arrangements might facilitate visual comparisons with Figure 4, where the ratios for the SR-based system and the IR-based system were plotted on the axes rising to the left and the right, respectively.

**Figure 6.**
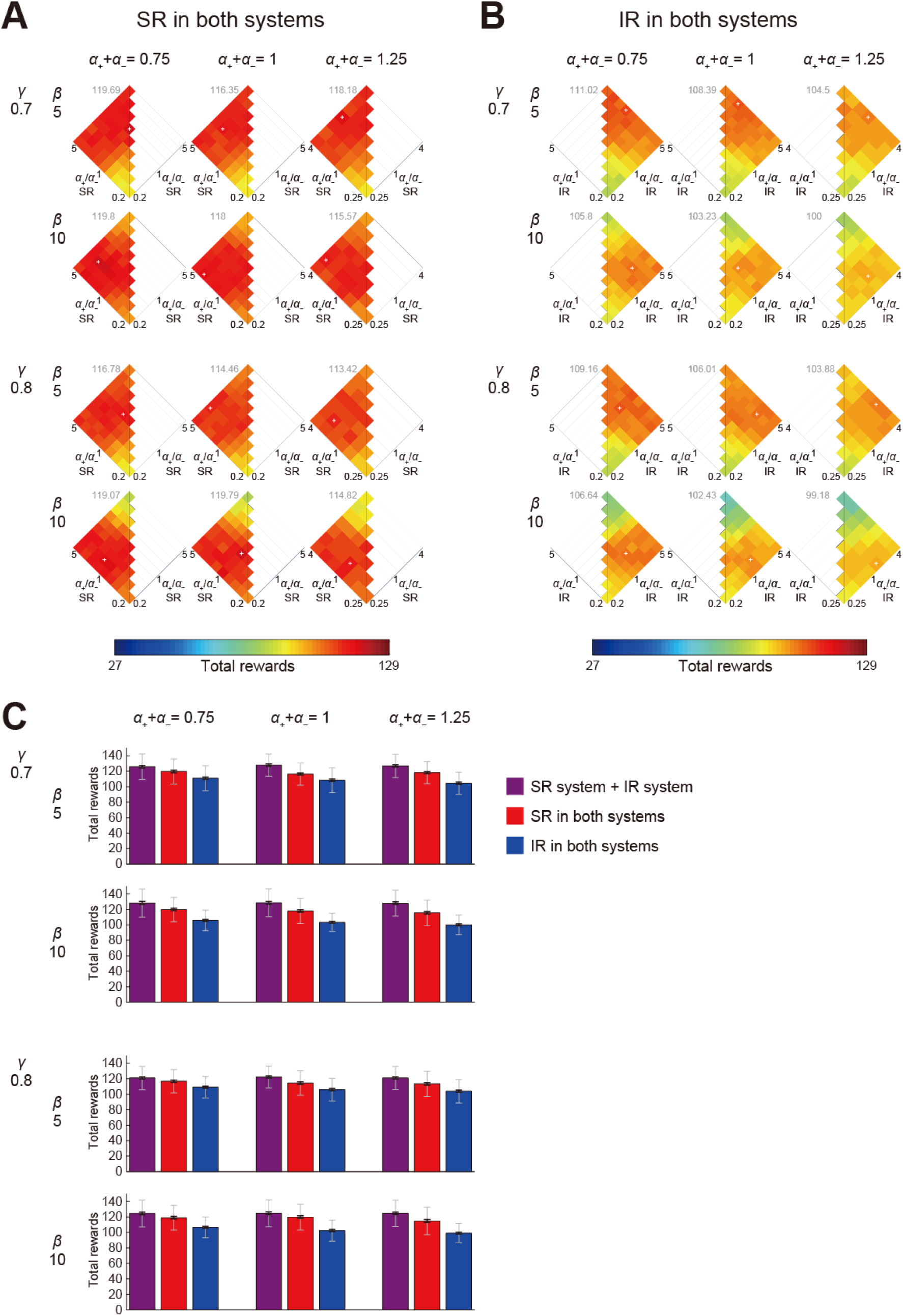
Performance of the model consisting of two systems, both of which employed the same way of state representation. Results in the case where both systems employed only the SR **(A)** or only the IR **(B)** are shown. Configurations are the same as those in Figure 4 (white cross: the best-performance set of *α*_+_/*α*_−_ ratios; gray number: the best performance (total rewards)). If the *α*_+_/*α*_−_ ratio was equal between both systems, the two systems behaved in exactly the same manner, and thus such conditions (on the vertical lines) were equivalent to having only a single system. Also, only the left (A) or right (B) side is shown, because “*α*_1+_ / *α*_1−_ = 0.2 and *α*_2+_ / *α*_2−_ = 3”, for example, are equivalent to “*α*_1+_ / *α*_1−_ = 3 and *α*_2+_ / *α*_2−_ = 0.2” given that both systems employed the same representation. The left (A)-and right-(B) placement was made so as to facilitate visual comparisons with Figure 4. **(C)** Comparison of the maximum performances (total rewards) of the cases where one system employed the SR while the other employed the IR (purple), both systems employed the SR (red), and both systems employed the IR (blue) for each set of the time discount factor (*γ*), inverse temperature (*β*), and the sum of learning rates for positive and negative TD-RPEs (*α*_+_ + *α*_−_). The bar lengths indicate the mean performance over 100 simulations for the condition that gave the best mean performance, and the black thick and gray thin error-bars indicate ±SEM and ±SD for that condition, respectively.

Comparing Figure 6A and 6B, SR-SR combination generally achieved better performance than IR-IR combination. This may not be surprising, given that a known feature of SR-based learning is sensitive adaptation to changes in rewards in the environments (Momennejad *et al*., 2017; Russek *et al*., 2017), which occurred in our simulated task. Besides, during the initial no-reward epoch, the SR-based system could acquire, through TD learning of SR features, an SR under the random policy, which presumably acted as beneficial “latent learning” (Dayan, 1993), while the IR-based system could not learn anything. Likewise, although the location of special reward candidate state and thus the optimal policy varied from epoch to epoch and SR is a policy-dependent representation, SR features acquired and updated in the agent should contain information about the basic structure of the grid space, which could help reward navigation throughout the task. Next, whether having two systems with different ratios of positive and negative TD-RPE-based learning rates was advantageous, in terms of the mean performance in this task, appears to be not clear in the cases where both systems employed only the SR or IR than in the case of the combination of SR- and IR-based systems. Last and most strikingly, the best combination of the SR-based and IR-based systems, i.e., the combination of the SR-based system learning mainly from positive TD-RPEs and the IR-based system learning mainly from negative TD-RPEs outperformed any combinations of two SR-based systems or two IR-based systems for all the sets of parameters (*α*_+_ + *α*_−_, *β*, and *γ*) that were so far examined (compare Figure 4 and Figure 6A,B; also shown in Figure 6C). We have conducted more comprehensive examination of the parameter space, and confirmed that the superiority of the combination of appetitive SR- and aversive IR-based systems still largely holds (see the Supplementary text, Figures S1 and S2, and Tables S1 and S2).

### Dependence on task properties

So far we examined the agent’s performance in the particular task. We examined how it could differ if task properties change. The original task contained a stochastic nature in a sense that reward was placed at a particular state (named the special reward candidate state) with a high probability (60%) in a given epoch but with the remaining probability reward was placed at one of the other (normal) candidate states. We examined what happened when this stochasticity was reduced or removed in the case of the model consisting of SR-based and IR-based systems with the original set of parameters used in Figures 3 and 5 (*α*_SR+_ + *α*_SR−_ = *α*_IR+_ + *α*_IR−_ = 1, *β* = 5, and *γ* = 0.7). Figure 7A shows the results for the cases where the probability of reward placement at the special candidate state was varied from 70% to 100%. As shown in the figure, as the stochasticity reduced, combinations of *α*_SR+_ / *α*_SR−_ and *α*_IR+_ / *α*_IR−_ that gave good performance gradually shifted, and when the stochasticity was totally removed, best performance was achieved when both *α*_SR+_ / *α*_SR−_ and *α*_IR+_ / *α*_IR−_ were high values.

**Figure 7.**
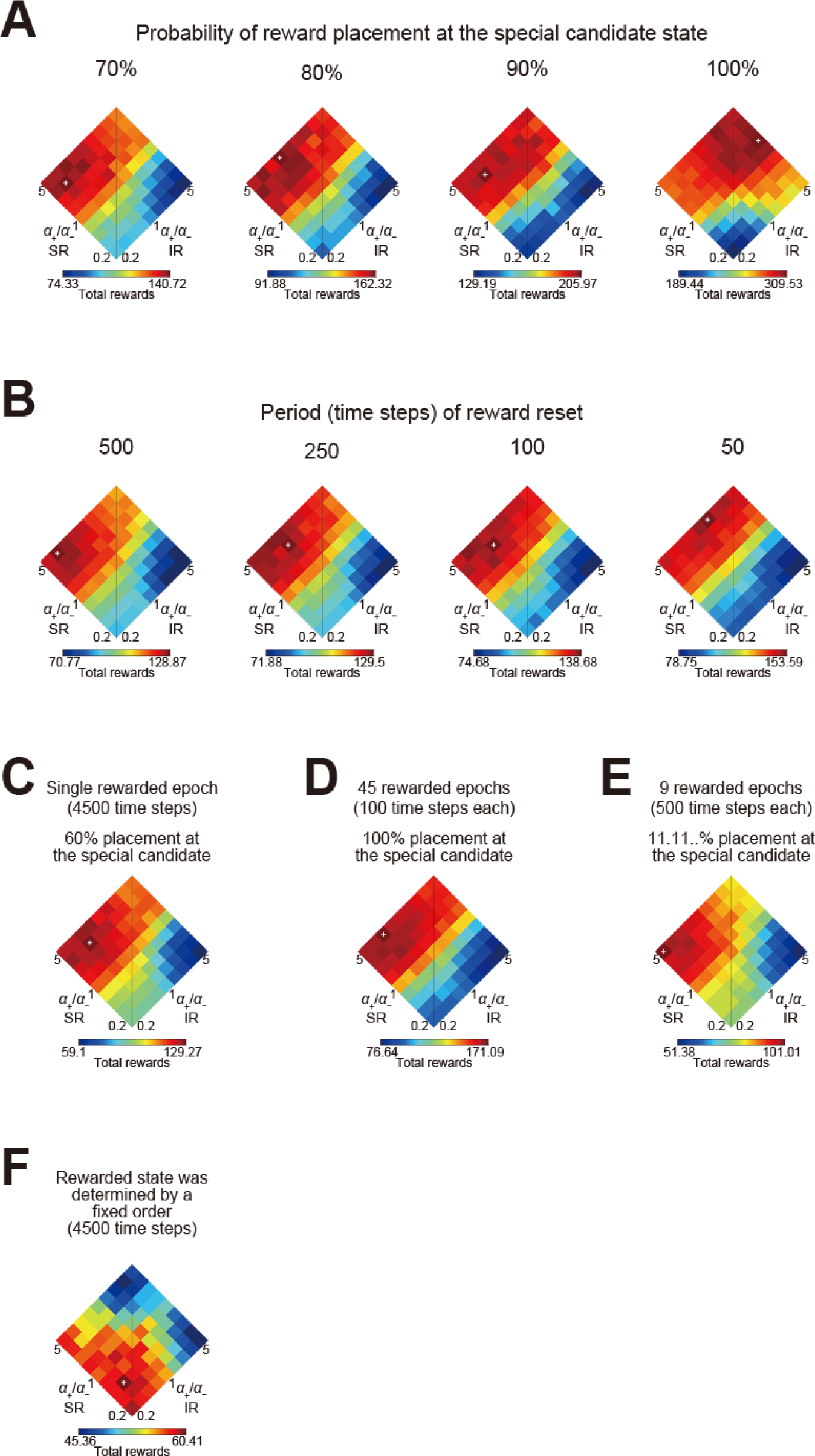
Performance of the model consisting of SR-based and IR-based system when task properties were changed. The model with the original set of parameters (*α*_SR+_ + *α*_SR−_ = *α*_IR+_ + *α*_IR−_ = 1, *β* = 5, and *γ* = 0.7) was used. The ranges of the color bars in this figure correspond to the ranges between the lowest and highest performances (i.e., the minimum and maximum total rewards) in the individual panels, and they are different from the color bars in the previous figures. **(A)** Performance (total rewards) for the cases where the probability of reward placement at the special candidate state was varied from 70% (leftmost panel), 80%, 90%, or 100% (rightmost). **(B)** Performance for the cases where reward location (state) was reset at every 500 (leftmost panel), 250, 100, or 50 (rightmost) time steps in the rewarded epochs; where reward was located was determined according to the original stochastic rule, i.e., reward was placed at the special candidate state for the epoch with 60% and placed at one of the other (normal) candidate states with 5% each. **(C)** Performance for the case where there was only a single rewarded epoch with 4500 time steps following the initial 500 time steps no-reward epoch. **(D)** Performance for the case where stochasticity of reward placement within each epoch was removed and the duration of each rewarded epoch was shortened from 500 time steps to 100 time steps while number of rewarded epochs was increased from 9 to 45. **(E)** Performance for the case where special candidate state was abolished and reward was placed at each of the nine candidate states with equal probability (1/9 = 11.11..%). **(F)** Performance for the case where rewarded state was determined by a fixed order, namely, (5, 1), (5, 5), (1, 5), and again (5, 1), and these were repeated throughout the task.

We also examined the effects of another variation in the task properties. In the original task, new reward was not placed until the agent obtained the previous reward. This property, coupled with the stochastic placement of reward, is considered to make perseveration-like behavior quite maladaptive and increase the importance of learning from negative feedbacks. We examined what happened if this property was weakened by introducing periodic resets of reward placement into the original task, again in the case of the model consisting of SR-based and IR-based systems with the original set of parameters. Figure 7B shows the results for the cases where reward location (state) was reset at every 500, 250, 100, or 50 time steps in the rewarded epochs; at the reset timings, reward was placed at the special candidate state for the epoch with 60% and placed at one of the other (normal) candidate states with 5% each. As shown in the figure, as the resets became more frequent, combinations of *α*_SR+_ / *α*_SR−_ and *α*_IR+_ / *α*_IR−_ that gave good performance again gradually shifted from high *α*_SR+_ / *α*_SR−_ and low *α*_IR+_ / *α*_IR−_to high *α*_SR+_ / *α*_SR−_ and also high *α*_IR+_ / *α*_IR−_, while overall sensitivity to *α*_IR+_ / *α*_IR−_ looked diminished.

We further examined three different variations of the original task. The first variation was abolishment of multiple rewarded epochs with different placements of special reward candidate state. Specifically, we examined the case where there was only a single rewarded epoch with 4500 time steps, instead of nine epochs with 500 time steps for each, following the initial 500 time-steps no-reward epoch. The special candidate state was varied only across simulations. The second variation was removal of the stochasticity of reward placement within each epoch, together with shortening of the duration of each rewarded epoch (from 500 time steps to 100 time steps) and increasing the number of rewarded epochs (from 9 to 45). The third variation was abolishment of special reward candidate state. Specifically, we came back to the original nine 500 time-steps rewarded epochs, but the probability of reward placement at the special candidate state was changed from the original 60% to 1/9 (11.11..%) so that reward was placed at each of the nine candidate states with equal probability (1/9). Figures 7C-E show the performance of the model consisting of SR-based and IR-based systems with the original set of parameters in these three variations. As shown in the figures, the patterns look similar to the case of the original task (Figure 3).

It was rather surprising that similar pattern appeared even in the last case, because reward was placed at any candidate states with equal probabilities and so it was no longer a task where the agent could learn where the special candidate state was in each epoch. However, the nine candidate states were neighboring with each other on the two edges of the grid space, and thus it would still have been the case that next reward was placed at a state near from the previously rewarded state with a relatively high probability and the agent could exploit it through positive TD-RPE-based learning. As a control, we examined a task in which positive TD-RPE-based learning was expected to have little meaning. Specifically, after 500 time-steps no-reward epoch, rewarded state was determined by a fixed order, namely, (5, 1), (5, 5), (1, 5), and again (5, 1), and these were repeated throughout the task. Figure 7F shows the performance of the model consisting of SR-based and IR-based systems with the original parameters. As shown in the figure, best performance, though rather low as expected, was achieved when both *α*_SR+_ / *α*_SR−_ and *α*_IR+_ / *α*_IR−_ were low values. This is reasonable because only quick erasing of the memory of previous reward is considered to be adaptive in this control task.

These results together indicate that the combination of appetitive SR and aversive IR systems does not always achieve the best performance, but does so in certain environments where reward placement is dynamically changing but still learnable using both positive and negative TD-RPEs.

### Performance of the two-system models in the two-stage tasks

Whether humans or animals take model-based/goal-directed or model-free behavior has been widely examined by using the so-called two-stage (or two-step) tasks (Daw et al., 2011; Groman et al., 2019b). Therefore we also examined how our models consisting of two systems performed in such tasks. We simulated the two-stage task (Daw *et al*., 2011). At the first stage, there were two choice options. Selection of each of these two options lead to one of two pairs of second-stage options with fixed probabilities (70% or 30%) (Figure 8A). Then, selection of one of the second-stage options lead to reward or no-reward outcome. The probability of reward for each second-stage option was independently set according to Gaussian random walk with reflecting boundaries at 0.25 and 0.75 (an example is shown in Figure 8Ba). The IR-based system learned the value of each first- and second-stage option through RPE-based updates. The SR-based system had the SR of the first- and second-stage options, and leaned their values through RPE-based updates of the weights of the approximate value function. The initial values of the SR matrix were set incorporating the probabilities of the transitions from the first stage to the second stage and also assuming random choice policy at the second stage, and the SR matrix was updated by using the prediction error of SR features.

**Figure 8.**
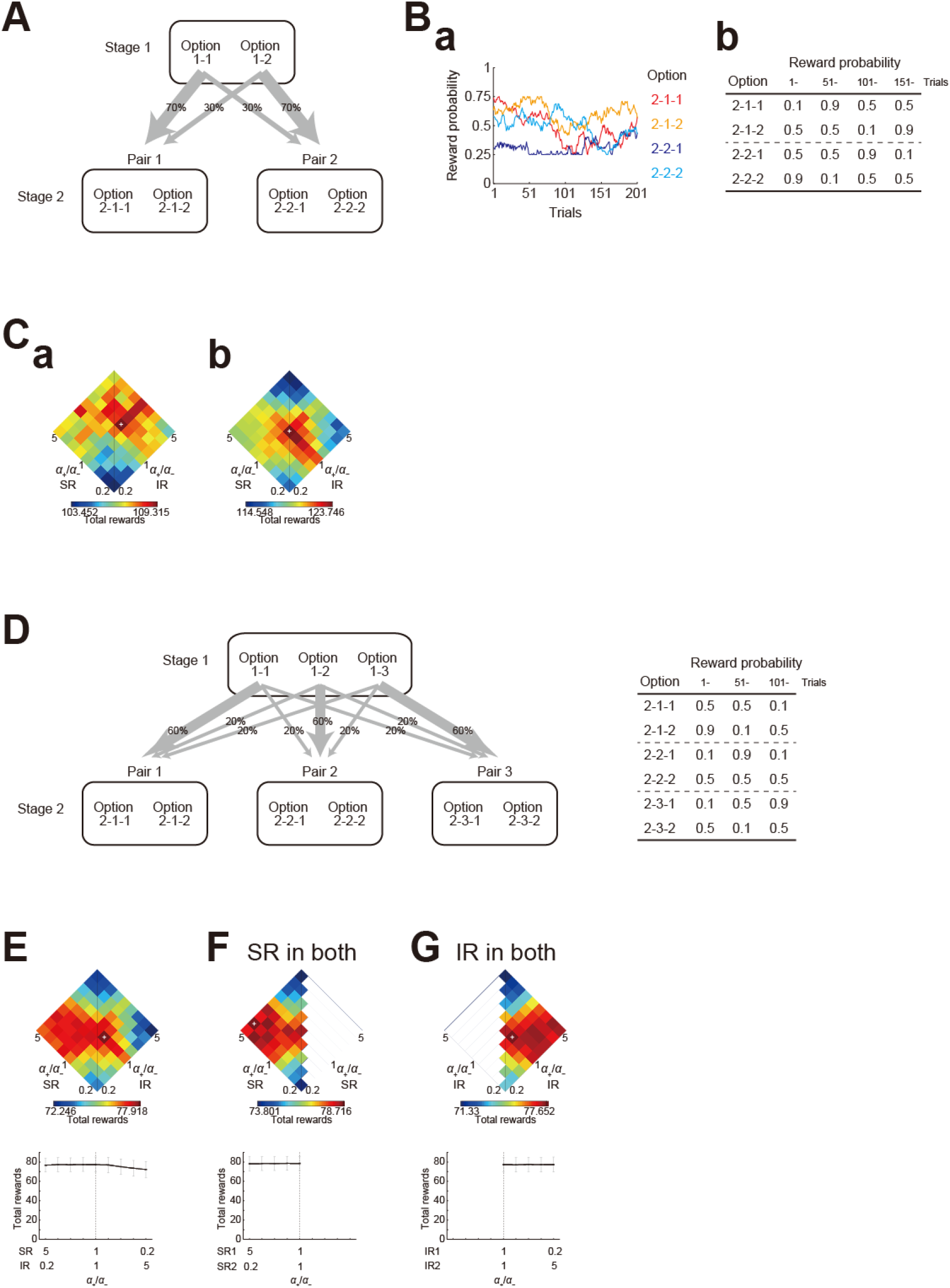
Performance of the two-system models in the two-stage tasks. **(A)** Schematic diagram of the two-stage task. Selection of one of the two first-stage options lead to one of the two pairs of second-stage options with fixed probabilities. **(B)** Reward probabilities for the four second-stage options. **(a)** In the original task, the probability of for each option was independently set according to Gaussian random walk with reflecting boundaries at 0.25 and 0.75. An example is shown. **(b)** In a variant of the task, the probabilities for the four options were set to specific values, which changed three times in the task. **(C)** Mean performance (mean total rewards over *n* = 1000 simulations) of the model consisting of an SR-based system and an IR-based system, with the ratio of positive- / negative-error-based learning rates (*α*_+_/*α*_−_) for each system was varied under *α*_+_ + *α*_−_ = 1, *β* = 5, and *γ* = 1. Panels (a) and (b) show the results for the tasks with the reward probabilities shown in (B)(a) and (B)(b), respectively. **(D)** *Left*: Schematic diagram of a variant of the two-stage task, in which there were three first-stage options and three pairs of second-stage options. *Right*: Reward probabilities for the six second-stage options, which were set to specific values and changed two times in the task. **(E-G)** *Top panels*: Mean performance of the model consisting of SR- and IR-based systems (E), two SR-based systems (F), or two IR-based systems (G), in the task variant with three first-stage options and three pairs of second-stage options. *Bottom graphs*: Mean (black solid line), SEM (black thick error-bars; though hardly visible), and SD (gray thin error-bars) of the performance over *n* = 1000 simulations for the conditions where *α*_+_/*α*_−_ for SR system times *α*_+_/*α*_−_ for IR system was equal to 1 (i.e., the conditions on the horizontal diagonal in the top panels).

Figure 8Ca shows the performance of the model with (*α*_+_ + *α*_−_ = 1, *β* = 5, and *γ* = 1) and the ratio of the learning rates for positive and negative RPEs in each system (*α*_SR+_/*α*_SR−_ and *α*_IR+_/*α*_IR−_) varied. As shown in the figure, combination of appetitive SR and aversive IR systems did not give a good performance in this task. We also examined a variant of the task, where the probabilities of reward for the second-stage options were set to specific values, which changed three times during the task (Figure 8Bb), so that differences among the values and their temporal changes could be clearer than the original case using random walk. Figure 8Cb shows the performance of the same model, and here again combination of appetitive SR and aversive IR systems did not give a good performance.

A possible reason why the good performance of such a combination observed in the navigation tasks was not generalized to the two-stage tasks was that the two-stage tasks imposed only selection from two options at both stages while the navigation tasks imposed selection from more than two options. In the case with only two choice options, an increase in the choice probability of one option simultaneously means the same magnitude of decrease in the choice probability of another option. Intuitively, generalization of learning from positive feedbacks for one option appears to have limited merit if choice probabilities are determined by the value difference between two options and there is no other option to which generalization can be applied. Therefore we simulated a variant of two-stage task, in which there were three, rather than two, first-stage options and three pairs of second-stage options (Figure 8D). Figure 8E shows the performance of the same model in this task. As shown in the figure, in this case, combination of appetitive SR-based system and aversive IR-based system achieved a relatively good performance. Conditions with similar intermediate *α*_+_/*α*_−_ ratios for both systems, or even with higher *α*_+_/*α*_−_ ratio for the IR system than for the SR system, achieved a good, or even better, mean performance, but shifts from such conditions towards appetitive IR- and aversive SR-based systems resulted in a rapid decrease in the performance while shifts towards appetitive SR- and aversive IR-based systems resulted in a milder decrease in the performance (although the overall range of the changes in the mean performance was rather small). Given this, we considered that the good performance of the combination of appetitive SR- and aversive IR-based systems observed in the navigation tasks could be generalized to the two-stage task with more than two options at least to a certain extent.

Figure 8F and G show the performance of the models in which both of the two systems employed only SR or IR, respectively. The ranges of the performance of these two types of models look comparable with each other, and also comparable to the performance range of the model consisting of SR- and IR-based systems. Therefore, the superiority of the solely SR-based model over the solely IR-based model, and also the superiority of the combined SR- and IR-based systems over the solely SR- (or IR-) based system, observed in the navigation tasks was not generalized to this two-stage task. It has been pointed out (Kool et al., 2016) that model-based control actually provides little performance benefits in the original two-stage task (Daw *et al*., 2011) as well as in several variants. Although model-based control can still be beneficial in certain two-stage tasks (Kool *et al*., 2016), advantages of model-based or SR-based control, and potentially also of the combined SR- and IR-based systems, might appear more clearly in tasks with more than two stages such as the navigation tasks.

### An extended model of cortico-basal ganglia circuits

In the cortico-basal ganglia circuits, it is likely that both the cortical neurons/regions using SR and those using IR target both the D1/direct and D2/indirect striatal projection neurons (SPNs), with potentially different weights. The targeted striatal regions, however, may differ depending on cortical sources. Also, although both D1 and D2 synapses could exhibit bidirectional plasticity (Shen et al., 2008), it is also possible that the D1/direct and D2/indirect pathways learn mostly from positive and negative TD-RPEs, respectively, given recent results (Lee *et al*., 2021) that phasic activation of DA neurons increased net activity of PKA in D1 SPNs while minimally modulating it in D2 SPNs whereas phasic inactivation of DA neurons increased net PKA activity in D2 SPNs but not significantly in D1 SPNs. In order to more accurately model these possibilities, we considered an extended model consisting of an SR-based system and an IR-based system, each of which consisted of two subsystems learning from either positive or negative TD-RPEs (modeling the D1/direct or D2/indirect pathway, respectively) (Figure 9A). To ensure convergence of the values of each subsystem, we introduced a decay/forgetting of the values (Mikhael and Bogacz, 2016) (see also (Morita and Kato, 2014) for value decay). We examined how this extended model performed in the original reward navigation task, varying the learning rate for each subsystem, specifically, *a*_SR+_ (for the subsystem learning from positive TD-RPEs of the SR-based system), *a*_SR−_, *a*_IR+_, and *a*_IR−_ under the condition with *a*_SR+_ + *a*_SR−_ = *a*_IR+_ + *a*_IR−_ = 1, *α*_SRfeature_ = 0.05, *β* = 5, *γ* = 0.7, and the decay rate per time step was 0.001. Notably, the learning rate *a*_SR+_, *a*_SR−_, *a*_IR+_, and *a*_IR−_ in this model can represent the strength of the connection between the corresponding cortical neurons/regions and striatal pathways (i.e., *a*_SR+_: from cortical neurons/regions using SR to the striatal direct pathway, *a*_SR−_: from those using SR to the indirect pathway, *a*_IR+_, and *a*_IR−_: from those using IR to the direct and indirect pathways; see the Materials and Methods).

**Figure 9.**
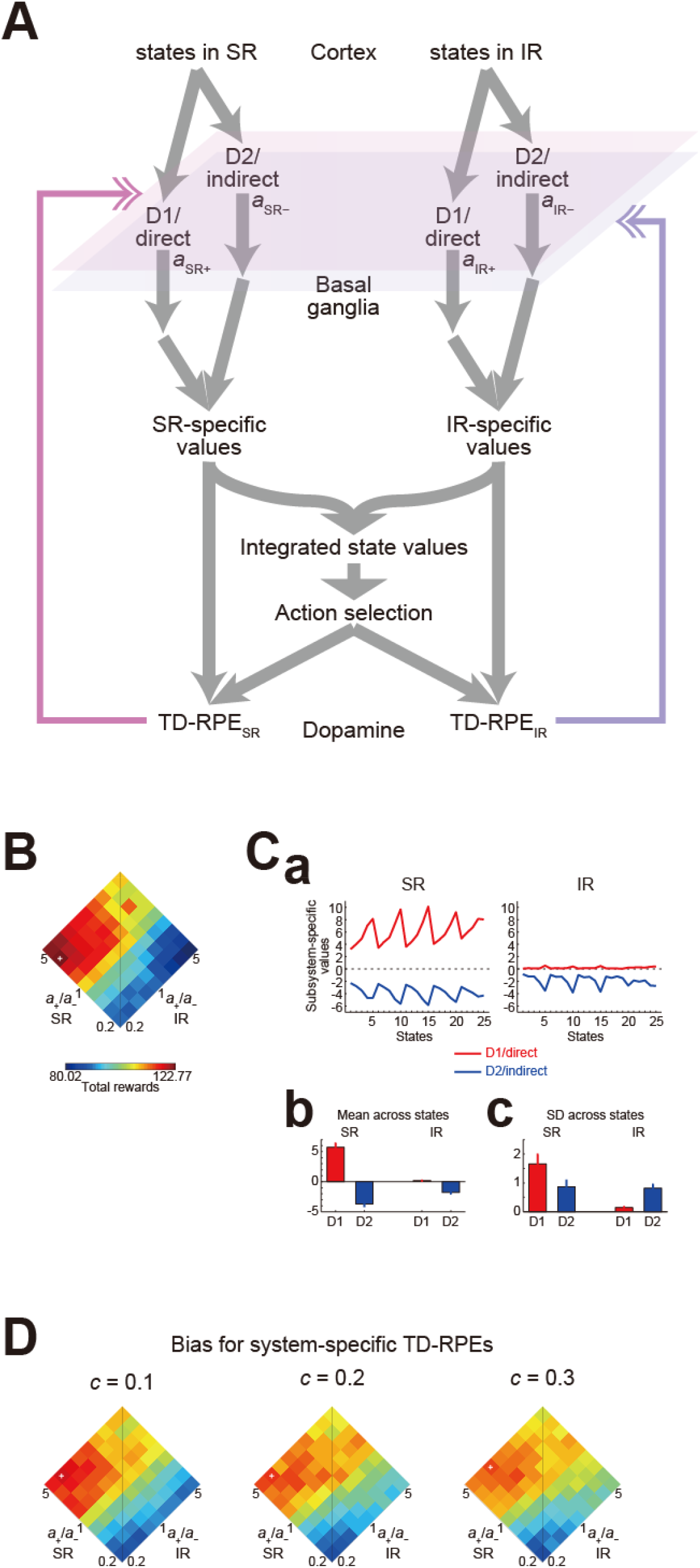
Performance of an extended model of cortico-basal ganglia circuits in the reward navigation task. **(A)** Schematic illustration of the model. The model consists of an SR-based system and an IR-based system, each of which consists of two subsystems learning from either positive or negative TD-RPEs with the learning rate *a*_SR+_ and *a*_IR+_ or *a*_SR−_ and *a*_IR−_, modeling the D1/direct or D2/indirect pathway of the basal ganglia, respectively. Action selection is based on the integrated state values, which are the means of the SR- and IR-system-specific values. SR- and IR-value-specific TD-RPEs are calculated, and they are received by each system without or with a bias for the system-value-specific TD-RPEs of the own system. **(B)** Mean performance in the cases without the bias for SR- and IR-value-specific TD-RPEs, plotting against the *a*_SR+_/*a*_SR−_ and *a*_IR+_/*a*_IR−_ ratios, under the condition with *a*_SR+_ + *a*_SR−_ = *a*_IR+_ + *a*_IR−_ = 1, *α*_SRfeature_ = 0.05, *β* = 5, *γ* = 0.7, and the rate of value decay per time step was 0.001. **(C)** Subsystem-specific values at the end of the task in the case of (*a*_SR+_, *a*_SR−_, *a*_IR+_, *a*_IR−_) = (0.8, 0.2, 0.2, 0.8). **(a)** Examples in a simulation. **(b**,**c)** Across-simulation means of the mean (b) and SD (c) of the subsystem-specific values across states. The error-bars indicate SDs across simulations. **(D)** Mean performance in the cases with the bias for SR- and IR-value-specific TD-RPEs, in which the SR-based system (both subsystems) receive (1 + *c*) times more SR-specific TD-RPEs than IR-specific TD-RPEs and vice versa. The color corresponds to the one in the color bar shown in (B).

Figure 9B shows the mean performance, plotting against the *a*_SR+_/*a*_SR−_ and *a*_IR+_/*a*_IR−_ ratios. As shown in the figure, combination of high *a*_SR+_/*a*_SR−_ and low *a*_IR+_/*a*_IR−_ ratios achieved good performance. Figure 9Ca shows examples of the subsystem-specific values at the end of the task in a simulation with (*a*_SR+_, *a*_SR−_, *a*_IR+_, *a*_IR−_) = (0.8, 0.2, 0.2, 0.8), and Figures 9Cb and 9Cc show across-simulation means of the mean and SD of the subsystem-specific values across states with the same set of parameters. As shown in the figure, the two subsystems, modeling the two basal ganglia pathways, developed antagonistic values in both the SR- and IR-based systems. Among the D1/direct-subsystems, the subsystem in the SR-based system had most information. On the other hand, the D2/indirect-subsystems in both SR- and IR-based systems had comparable information.

**Figure 10.**
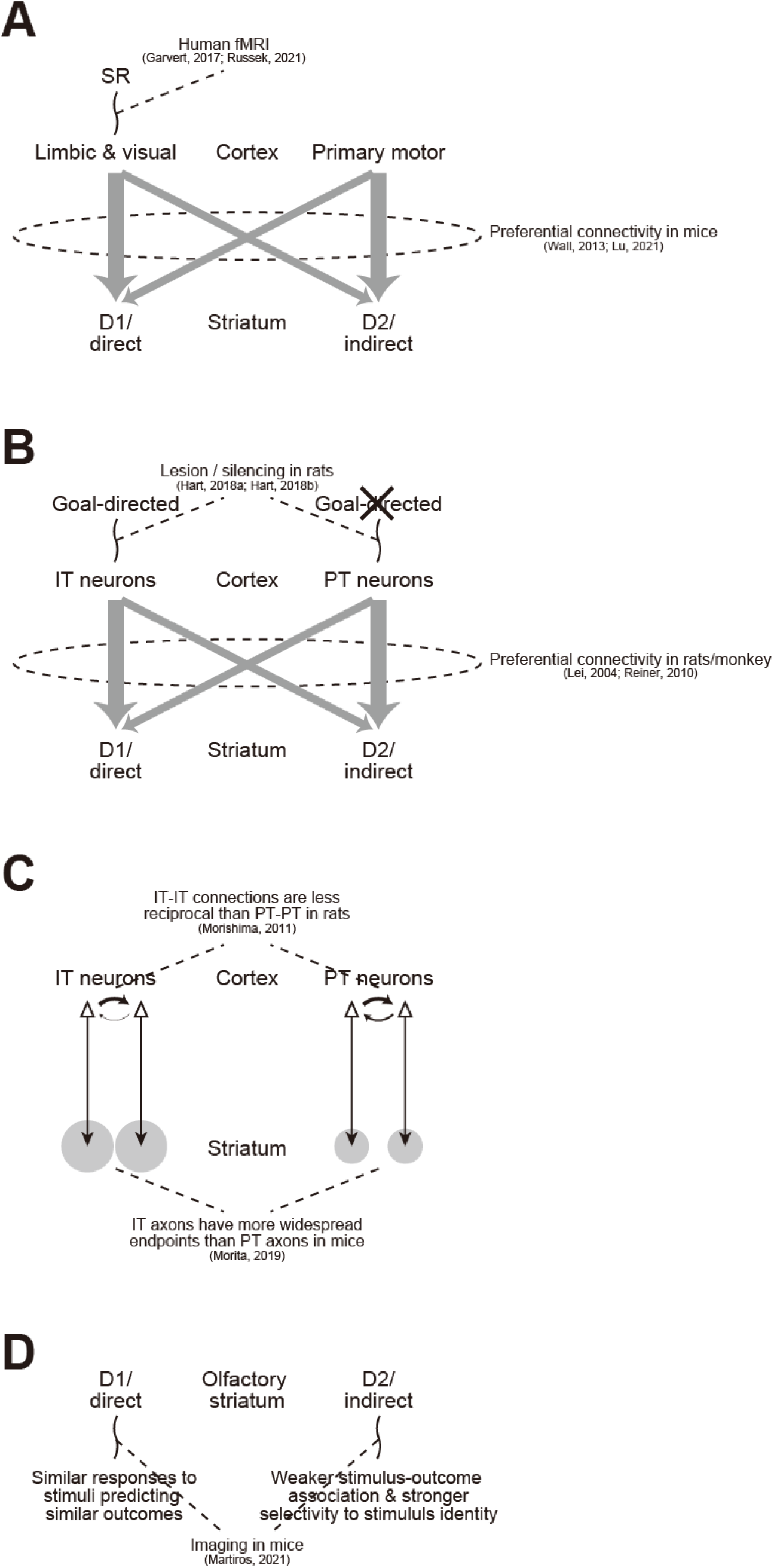
Explanation for the significance and mechanism of diverse findings about the cortico-BG circuits. **(A)** Experimentally suggested limbic/visual cortical encoding of SR and limbic/visual->D1/direct and primary motor->D2/indirect preferential connections indicate preferential use of SR in the appetitive D1/direct pathway, whose merit was explained by our model. **(B)** Experimentally suggested involvement of IT-type, but not PT-type, corticostriatal neurons in goal-directed behavior and IT->D1/direct and PT->D2/indirect preferential connections also indicate preferential use of SR in the appetitive D1/direct pathway, whose merit was explained by our model. **(C)** Less reciprocal IT-IT connections and wider IT->striatum axonal endpoints are in line with engagement of IT (rather than PT) neurons in SR-like representation. **(D)** D1 neurons’ similar responses to stimuli predicting similar outcomes and D2 neurons’ weaker stimulus-outcome association and stronger selectivity to stimulus identity in the ventral striatal olfactory tubercle could be explained by preferential use of SR- and IR-like representations in the D1 and D2 pathways, respectively, as adopted in the superior combination in our model.

Recently it was suggested that different parts of the striatum would receive inhomogeneous DA signals (Hamid et al., 2021) and also that differential DA signals in (relatively nearby) striatal regions may reflect representation-specific components of TD-RPEs (Lee et al., 2022). Given these, we also examined a case where the SR-based system and the IR-based system did not receive exactly the same TD-RPEs but received partially different signals. Specifically, we calculated SR-value-specific TD-RPEs and IR-value-specific TD-RPEs using SR- or IR-specific values, and assumed that the SR-based system (both subsystems) receive (1 + *c*) times more SR-specific TD-RPEs than IR-specific TD-RPEs and vice versa, with *c* was varied to 0.1, 0.2, or 0.3, while the sum of the TD-RPEs received by the SR- and IR-based systems was kept unchanged from the original case (corresponding to *c* = 0). Figure 9D show the mean performance of these model variants. As shown in the figure, the superiority of the combination of high *a*_SR+_/*a*_SR−_ and low *a*_IR+_/*a*_IR−_ ratios was largely preserved, while the achieved highest performance decreased as *c* increased.

## Discussion

We found that combination of SR-based system learning mainly from positive TD-RPEs and IR-based system learning mainly from negative TD-RPEs showed superior performance in certain dynamic reward environments. Below we discuss reasons for the superiority of such a combination, and show how such a combination can explain the functional significance and underlying mechanism of diverse findings about the cortico-BG circuits. We also discuss limitations and perspectives.

### Reasons for the superior performance of the combined appetitive SR- and aversive IR-based systems

As possible reasons why the combination of appetitive SR- and aversive IR-based systems performed well, the following two are considered. The first possible reason comes from an asymmetry between positive- and negative-error-based learning. When the agent unexpectedly encountered a reward at a certain state, next reward was placed at the same state or nearby states with a certain high probability (given that there was learnable structure). Then, quickly revising up the value of not only the rewarded state itself but also any states from which the rewarded state could be easily reached (i.e., states having high SR feature values) would be beneficial. In other words, SR-dependent generalization of positive TD-RPE-based value updates would be beneficial. The result that the combination of appetitive SR- and aversive IR-based systems performed relatively well in the two-stage task with three first-stage options but not in the task with two options seems in line with this, because a key difference between the task variant with two options only and the one with three options would be that generalization would have a larger merit in the latter than in the former. Next, assume that the agent obtained rewards several times at the same or nearby states (in the navigation task), and then reward was placed at a distant state. This time, the agent was likely to unexpectedly encounter reward omission. Critically, through the repetitive acquisition of rewards at the nearby states, the agent’s policy was likely to be sharpened and therefore revising down the values of only the states right on the sharpened policy would already be effective. In other words, as for negative TD-RPE-based value updates, IR-based narrow effect would be largely sufficient, and too much SR-dependent generalization could rather be harmful.

The second possible reason for the good performance of the combination of appetitive SR- and aversive IR-based systems, which is considered to be also related to the last point of the above, comes from the policy-dependence of SR. SR reflects state transitions under the policy that has been used. Thus, when reward is relocated at a distant place and thereby optimal policy is drastically changed, SR under the policy optimized for the previous reward should significantly differ from the one under the new optimal policy, and SR becomes updated as the agent changes its policy. In fact, through TD-RPE-based learning, in contrast to direct calculation of state values by multiplication of SR and reward at each state (if it could be estimated), value function after reward relocation can in principle be approximated even with SR under the previously near-optimal policy, as well as with arbitrary state representation, as suggested in (Russek *et al*., 2017). Nonetheless, SR under the previously near-optimal policy may not have fine information about state transitions near the new reward location, implying potential difficulty in learning from negative TD-RPEs for SR-based system, which could contribute to the superiority of the combination of appetitive SR- and aversive IR-based systems. Difficulty in the case with drastic change in the goal location and the optimal policy has previously been shown (Lehnert et al., 2017) for learning using successor features, which are generalization of SR (Barreto et al., 2016). Increasing the learning rate for SR feature update (*α*_SRfeature_) could potentially mitigate this issue. In our simulations with *α*_SRfeature_ varied (Figure S1 and Table S1), as *α*_SRfeature_ increased, combination of the *α*_+_/*α*_−_ ratios in the two systems that achieved good performance approached to the ones with similar ratios in both systems, in line with the above argument. At the same time, however, increase in *α*_SRfeature_ did not generally improve the highest achievable performance of the model with SR- and IR-based systems; increase up to 0.1 improved it in some but not other conditions, and further increase could rather worsen the highest achievable performance presumably because state representation became too unstable, while increase in *α*_SRfeature_ up to 0.15 or 2 was beneficial in many cases for the model consisting of two SR-based systems (Figure S2 and Table S1). From the results that relatively small *α*_SRfeature_ was good for the model with appetitive SR- and aversive IR-based systems, we speculate that policy-independent state representation based on the transition structure, specifically, default representation (Piray and Daw, 2021), could alternatively be used.

### Explanation for the significance and mechanism of diverse findings about the cortico-BG circuits

The superiority of the combination of SR-based appetitive and IR-based aversive learners shown in our model provides a novel coherent explanation for the functional significance and underlying mechanism of diverse findings about the cortico-BG circuits, which could not be explained by previous dual-systems models that did not consider different representations in the D1/direct and D2/indirect pathways.

First, monosynaptic rabies virus tracing in mice revealed preferential connections from the limbic/visual cortices and primary motor cortex to the D1/direct and D2/indirect pathways, respectively (Lu *et al*., 2021; Wall *et al*., 2013), while human fMRI experiments found activations indicative of SR in the limbic/visual cortices, or more specifically, hippocampal–entorhinal cortex (Garvert *et al*., 2017) and visual cortex (Russek *et al*., 2021) (Figure 10A). These findings together indicate preferential use of SR in the D1/direct pathway (and not in the D2/indirect pathway), and the results of our simulations explain a functional merit of such a combination.

Second, electron microscopic analyses focusing on synapse sizes in rats (Lei *et al*., 2004) and monkey (Reiner *et al*., 2010) indicated preferential connections from the intratelencephalic (IT)- and pyramidal-tract (PT)-type corticostriatal neurons to the D1/direct and D2/indirect pathways, respectively, while lesion and silencing experiments in rats (Hart et al., 2018a; Hart et al., 2018b) demonstrated that prelimbic IT neurons, but not PT neurons, mediated sensitivity to outcome devaluation (Figure 10B), which is a defining feature of goal-directed behavior and can be achieved through SR-based learning (we have confirmed this: see the Supplementary text and Figure S3). These findings together are in line with preferential use of SR in the D1/direct pathway (and not in the D2/indirect pathway), and again, the results of our simulations explain a functional merit of such a combination.

Third, local connections between IT neurons were shown to be less reciprocal (i.e., more unidirectional) than those between PT neurons in rat frontal cortex (Morishima et al., 2011), and analysis of the MouseLight database (Winnubst et al., 2019) revealed that axonal projections of individual IT neurons to the striatum have on average more widespread endpoints than those of PT neurons (Morita et al., 2019) (Figure 10C). These properties are also in line with the possibility that IT neurons (rather than PT neurons) are engaged in SR-like representation, because SR is based on the directional transition relationships between states and also calculation of state value using SR requires access to not only that state but also all the states which are reached from that state through transitions.

Last but not least, recent work examining the ventral striatal olfactory tubercle of mice (Martiros et al., 2021) found that D1 neurons showed similar responses to stimuli predicting similar outcomes whereas D2 neurons showed weaker stimulus-outcome association and stronger selectivity to stimulus identity (Figure 10D). These differential responses could be explained by preferential use of SR- and IR-like representations in the D1 and D2 pathways, respectively, as adopted in the superior combination in our model, although the olfactory tubercle is known to receive direct inputs from the olfactory bulb (Igarashi et al., 2012) and it seems unclear whether there exist region/neuron-type- and pathway-dependent connection preferences homologous to those suggested for other (more dorsal) striatal region.

### Limitations and perspectives

As discussed above, our dual systems model with different representations potentially well describes neuron type- and region-specific circuitries in the cortico-BG system, going beyond the previous models that took into account the two BG pathways but not the diversity of the neocortex. However, there are many limitations. Sets of parameters that we examined were still limited. It also remains to be seen whether the superiority of the combination of appetitive SR-based and aversive IR-based systems is applied also to other tasks, including more complex tasks. Neurobiologically, while we considered appetitive and aversive learning in the two BG pathways, there are studies suggesting that the roles of these pathways differ not merely in the valence (Iino *et al*., 2020; Matamales et al., 2020; Nonomura et al., 2018; Peak et al., 2020). Also, mechanisms of TD-RPE calculation in DA neurons (Morita and Kawaguchi, 2019; Watabe-Uchida et al., 2017), as well as richer temporal dynamics and functional roles of DA suggested by recent studies (Bogacz, 2020; Hamid *et al*., 2021; Mikhael et al., 2022), remain to be incorporated.

A central prediction of our present modeling work is that humans and other animals take the learning strategy that was suggested to be superior by the model, i.e., combination of SR-based appetitive and IR-based aversive learning. This can in principle be tested by examining whether the behavior exhibits features of SR-based learning (c.f., (Momennejad *et al*., 2017)) differently between the cases with positive and negative feedbacks, potentially also with neuroimaging (c.f., (Garvert *et al*., 2017; Russek *et al*., 2021)).

## Materials and Methods

### Simulated reward navigation task

We simulated a reward navigation task in a certain dynamic reward environment. An agent was moving around in a 5 × 5 grid space (Figure 1A). The agent started from the fixed start state, which was location (1, 1), and moved to one of the neighboring states (two states at the corners, three states at the edges, and four states elsewhere) at each time step. During the initial 500 time steps, there was no reward, and the agent just moved around (Figure 1B). After that, a reward (always size 1) was introduced into one of the state (location) in the space. There were nine reward candidate states, where reward could potentially be placed, namely, (1, 5), (2, 5), (3, 5), (4, 5), (5, 1), (5, 2), (5, 3), (5, 4), and (5, 5) (i.e., the states on the two edges apart from the start state). During the next 500 time steps following the initial no-reward epoch, one of these nine candidate states was specified as the *special reward candidate state*, whereas the remaining eight candidate states were regarded as *normal candidate states*. There were in total nine rewarded epochs, each of which lasted for 500 time steps, and each one of the nine candidate states became the special candidate state in one of the nine epochs; the order was determined by pseudorandom permutation in each single simulation.

In the rewarded epochs, if the agent reached the rewarded state and obtained the reward, the agent was immediately carried back to the start state, and a new reward was introduced into a state, which was the special reward candidate state with 60% probability and one of the eight normal reward candidate states with an equal probability (i.e., 5%) each (Figure 1C).

We also examined variations of the task. In all the variations, there was the initial 500 time-steps no-reward epoch. In the first variation, the probability that reward was placed at the special candidate state, which was 60% in the original task, was varied to 70, 80, 90, or 100%. In the second variation, periodic resets of reward placement were introduced. Specifically, reward location (state) was reset at every 500, 250, 100, or 50 time steps in the rewarded epochs; at the reset timings, reward was placed at the special candidate state for the epoch with 60% and placed at one of the other (normal) candidate states with 5% each. In the third variation, the original nine rewarded epochs with different placements of special reward candidate state (500 time steps each) were abolished and instead there was only a single rewarded epoch with 4500 time steps. For this task variation, the special candidate state was varied only across simulations. In the fourth variation, reward was always placed at the special candidate state in each epoch and the duration of rewarded epoch was shortened from 500 time steps to 100 time steps while the number of rewarded epochs was increased from 9 to 45. The order of special reward candidate states was determined by five consecutive pseudorandom permutations of the nine candidate states. In the fifth variation, the probability of reward placement at the special candidate state was changed to 1/9 (11.11..%) so that reward was placed at each of the nine candidate states with equal probability (1/9). In the sixth variation, rewarded state was determined by a fixed order, namely, (5, 1), (5, 5), (1, 5), and again (5, 1), and these were repeated throughout the task.

### Model agent equipped with two learning systems

The agent’s behavior was controlled by an RL model consisting of two learning systems (Figure 2). The two systems may use different ways of state representation. We considered the successor representation (SR) and the individual representation (IR) (also called the “punctate” representation), and examined the cases where system 1 and 2 used the SR and IR, respectively, and also the cases where only the SR or IR was used in both systems. Each system had its own *system-specific value* of each state (detailed below). The mean (average) of the system-specific values of the two systems was calculated and used as the *integrated value* of each state *S*, denoted as *V*(*S*). At any states other than the rewarded state, the agent selected an action to move to one of the neighboring states depending on their integrated state values in a soft-max manner. Specifically, the agent selected the action to move to state *S*_*i*_, among the neighboring states *S*_*j*_, with probability

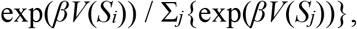

where *β* indicates the inverse temperature parameter, representing the degree of exploitation over exploration. When the agent was at state *S*(*t*) at time-step *t* and moved to *S*(*t*+1) at *t*+1, where *S*(*t*) was not the rewarded state, TD-RPE was calculated based on the integrated state values:

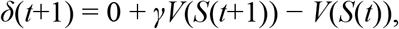

where *γ* indicates the time discount factor. When the agent was at *S*(*t*), which was the rewarded state, TD-RPE

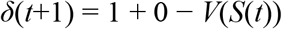

was calculated, again using the integrated state value.

In the system using the IR, the system-specific state value, *V*_*system*-*i*_(*S*(*t*)), where *i* was 1 or 2, was directly updated based on the TD-RPE *δ*(*t*+1), with potentially different learning rates depending on whether the TD-RPE was positive (non-negative) or negative:

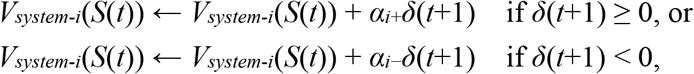

where *α*_*i*+_ and *α*_*i*−_ indicate the learning rates of the system *i* for positive (non-negative) and negative TD-RPEs, respectively, and they were systematically varied as described in the Results. The initial values of the system-specific state values of the system using the IR were set to 0.

In the system using the SR, the system-specific state value, *V*_*system*-*i*_(*S*(*t*)), where *i* was 1 or 2, was calculated as a linear function of the feature variables of the state. In the SR, every state *S*_*i*_ was represented by a set of feature variables, which were estimated cumulative discounted future occupancies of all the states *S*_*j*_ (*j* = 1, …, 25), denoted as *σ*(*S*_*i*_, *S*_*j*_). These feature variables were updated through TD learning of SR features (Gardner et al., 2018; Gershman et al., 2012). Specifically, when the agent was at *S*(*t*) at *t* and moved to *S*(*t*+1) at *t*+1, where *S*(*t*) was not the rewarded state, the TD errors for SR features were calculated as:

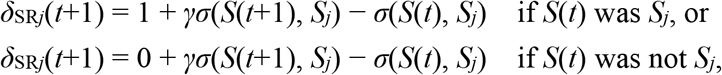

for all the states *S*_*j*_ (*j* = 1, …, 25). If *S*(*t*) was the rewarded state, the term *γσ*(*S*(*t*+1), *S*_*j*_) was dropped. Using these TD errors, *σ*(*S*(*t*), *S*_*j*_) (*j* = 1, …, 25) were updated as:

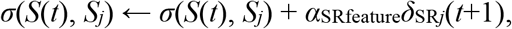

where *α*_SRfeature_ indicates the learning rate for the update of SR features and was set to 0.05 unless otherwise mentioned; *α*_SRfeature_ = 0.1, 0.15, 0.2, and 0.25 were also examined in Figures S1 and S2 / Tables S1 and S2. The initial values of all the feature variables were set to 0. The system-specific state value, *V*_*system*-*i*_(*S*(*t*)), was given as a linear function of the feature variables as:

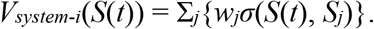

The weights (coefficients) *w*_*j*_ (*j* = 1, …, 25) were updated based on the TD-RPE *δ*(*t*+1) described above, with potentially different learning rates depending on whether the TD-RPE was positive (non-negative) or negative:

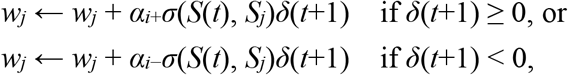

where *α*_*i*+_ and *α*_*i*−_ indicate the learning rates of the system *i* for positive (non-negative) and negative TD-RPEs, respectively, and they were systematically varied as described in the Results. The initial values of the weights *w*_*j*_ were set to 0.

### Simulated two-stage tasks

We also simulated the two-stage task (Daw *et al*., 2011) and its variants. In our simulation of the original task (Figure 8A), there were two choice options at the first stage. Selection of one of them led to each of two pairs of second-stage options with fixed probabilities (70% or 30%), whereas selection of the other first-stage option led to each pair of second-stage options with the opposite probabilities (i.e., 30% and 70%). Then, selection of one of the second-stage options led to reward or no-reward outcome. The probability of reward for each second-stage option was independently set according to Gaussian random walk with reflecting boundaries at 0.25 and 0.75. More specifically, the reward probabilities for the four second-stage options were independently changed at each trial by adding pseudo normal random numbers with mean 0 and SD 0.025, and reflected at 0.25 and 0.75, throughout the task consisting of 201 trials (Figure 8Ba).

We also simulated a variant of the task, where the probabilities of reward for the four second-stage options (two options for each of the two pairs) were set to specific values and the option-probability contingency was changed three times during the task consisting of 201 trials. Specifically, the probabilities were initially set to (0.1 and 0.5) for the first and second option of the first pair and (0.5 and 0.9) for the first and second option of the second pair, respectively, and changed to (0.9 and 0.5) and (0.5 and 0.1) at the 51th trial, (0.5 and 0.1) and (0.9 and 0.5) at the 101th trial, and (0.5 and 0.9) and (0.1 and 0.5) at the 151th trial (Figure 8Bb).

We further simulated another variant of the task, in which there were three, rather than two, first-stage options and three pairs of second-stage options (Figure 8D, left). Selection of each of the three first-stage options lead to one of three pairs of second-stage options with fixed probabilities: (60%, 20%, 20%), (20%, 60%, 20%), and (20%, 20%, 60%). The probabilities of reward for the six second-stage options (two options for each of the three pairs) were set to specific values and the option-probability contingency was changed two times during the task consisting of 150 trials. Specifically, the probabilities were initially set to (0.5, 0.9), (0.1, 0.5), and (0.1, 0.5) for the (first, second) option of the first, second, and third pair, and changed to (0.5, 0.1), (0.9, 0.5), and (0.5, 0.1) at the 51th trial, and (0.1, 0.5), (0.1, 0.5), and (0.5, 0.9) at the 101th trial (Figure 8D, right).

In all the variants of the two-stage tasks, at every trial, SARSA-type TD RPE for the first stage was calculated after the second-stage choice was determined:

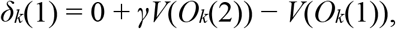

where *O*_*k*_(1) and *O*_*k*_(2) represent the chosen option for the first and second stage at the *k*-th trial, respectively. *V*(*O*) represents the value of option *O. γ* is the time discount factor, which was assumed to be 1 (i.e., no temporal discounting) in the two-stage tasks. Then TD RPE for the second stage:

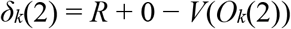

was calculated after reward (*R* = 1 or 0) was determined, where *V*(*O*_*k*_(2)) reflected *δ*_*k*_(1)-based updates of the weights of the approximate value function (see below) in the cases with SR-based system(s). The IR-based system(s) learned the value of each first- and second-stage option (in total 2+4=6 options or 3+6=9 options) through RPE-based updates, with the initial values set to 0. The SR-based system(s) had the SR of the first- and second-stage options, and leaned their values through RPE-based updates of the weights of the approximate value function. The SR matrix (6×6 or 9×9) was initialized to the one under the random policy regarding the choice at the second stage, incorporating the presumed stage-transition probabilities. The SR matrix was then updated by using the prediction errors of SR features. More specifically, at every trial, the SR features for the option chosen at the first stage was updated according to the prediction errors after the stage transition occurred and the second-stage option was determined, with the learning rate set to 0.05. Choice at both stages was made in the soft-max manner with the degree of exploitation over exploration (inverse temperature) (*β*) set to 5.

### Simulated outcome devaluation test

In a separate set of simulations, we simulated an outcome devaluation test. For this, we considered a narrowed grid space (Figure S3A). There were two goal states, Goal 1 at (3, 1) and Goal 2 at (1, 3), and when the agent reached one of the pre-goal states, i.e., (2, 1) or (1, 2), in the next time step it always proceeded to the corresponding goal state unless it was at the end of task epoch. There was an initial 500 time-steps no-reward epoch (Figure S3B), in which the agent was carried back to (1, 1) when it reached either goal state. Following the no-reward epoch, a 500 time-steps learning epoch started, and two rewards were introduced at the two goal states. During the learning epoch, when the agent reached either of the goal states, it obtained the reward, and reward was immediately resupplied. At the beginning of the learning epoch, there was a forced learning period, in which the agent was alternately placed at either pre-goal state, (2, 1) or (1, 2), with its order counterbalanced across simulations, in every 2 time steps for 10 times for each pre-goal state so that it forcedly experienced each reward 10 times. After the forced learning period has ended, the agent was carried back to (1, 1), and thereafter it freely behaved according to learned state values; when it reached either goal state, it was carried back to (1, 1).

After the learning epoch has ended, there was a devaluation phase, in which the agent experienced devaluation of one of the rewards, specifically, the reward introduced at Goal 2 (1, 3). The agent was directly placed at this goal state, and experienced TD error-based updates, but did not experience state transition. After this devaluation phase, agent was placed at (1, 1), and which of the two pre-goal states the agent initially reached was examined.

We examined the behavior of agent equipped with combination of SR-based and IR-based systems, with the ratios of the learning rates from positive and negative TD-RPEs in the two systems (*α*_SR+_/*α*_SR−_ and *α*_IR+_/*α*_IR−_) varied while their multiplication, as well as the sum of positive- and negative TD-RPE-based learning rates, were kept constant at 1 (*α*_SR+_/*α*_SR−_ × *α*_IR+_/*α*_IR−_ = 1 and *α*_SR+_ + *α*_SR−_ = *α*_IR+_ + *α*_IR−_ = 1; i.e., conditions corresponding to those on the horizontal diagonal in Figure 3A). As a control, we also examined the behavior of agent equipped with two IR-based systems, with the ratios of the learning rates from positive and negative TD-RPEs in the two systems similarly varied. Learnings of system-specific values, weights of the value function, and SR features were all done in the same manner as in the case of the reward navigation task, with the parameters *β, γ*, and *α*_SRfeature_ were set to 5, 0.7, and 0.05 (the same as those used for Figure 3); *β* = 10 was also examined in Figure S3Db. For each condition, the proportion of reaching the pre-goal state preceding the devalued Goal 2 in the test period in 100 simulations was calculated for 50 sets (i.e., in total 5000 simulations were made for each condition), and its mean and standard deviation were plotted for each condition in Figure S3D.

Notably, the assumed automatic transition from pre-goal state to goal state was critical. In our model, agent chose an action to move to one of the neighboring states according to their state values. The state values can thus be equated with the action values, or action policy, at the preceding states. Therefore, reduction in the value of goal state due to reward devaluation, which occurs even in the case of purely IR-based agent (i.e., agent with two IR-based systems), simultaneously means that the “action value” of “action” from the preceding state to the goal state is reduced. By contrast, it is usually considered that the value of outcome and the value of action which leads to the outcome are not equated but distinguishable, and outcome devaluation decreases the former by definition but can spare the latter. In order to be consistent with this, we assumed the “pre-goal state”, and assumed that “action” from the pre-goal state to the goal state was automatic and its value was not distinguishable from the goal value while the value of action leading to the pre-goal state was distinguishable.

### An extended model of cortico-basal ganglia circuits

We considered an extended agent’s model consisting of an SR-based system and an IR-based system, each of which consisted of two subsystems learning from either positive or negative TD-RPEs (modeling the direct or indirect pathway, respectively) (Figure 9A). The subsystems of the IR-based system learning from positive and negative TD-RPEs developed subsystem-specific values *V*_IR+_ and *V*_IR−_, respectively, according to the following rules:

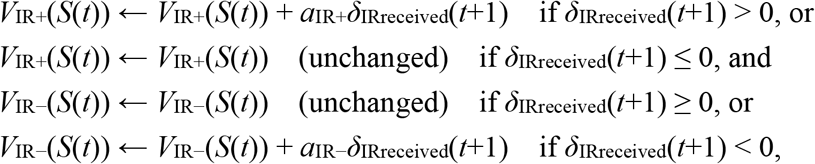

where *δ*_IRreceived_(*t*+1) represents the TD-RPE received by (both subsystems of) the IR-based system (see below), and also a constant decay/forgetting at every time-step:

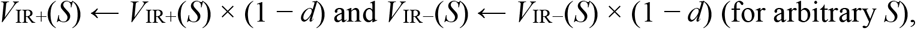

where *d* represents the rate of decay, which was set to 0.001. The IR-based system specific value *V*_IR_(*S*(*t*)) was calculated as the sum of *V*_IR+_(*S*(*t*)) and *V*_IR−_(*S*(*t*)):

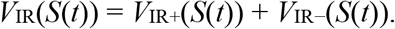

The subsystems of the SR-based system learning from positive and negative TD-RPEs developed subsystem-specific weights *w*_+*j*_ and *w*_−*j*_ (*j* = 1, …, 25), respectively, according to the following rules:

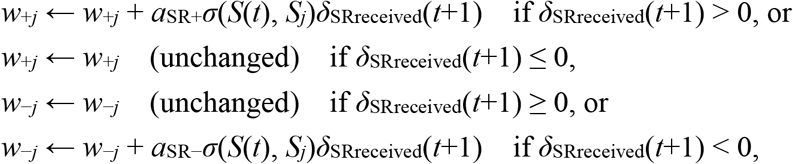

where *δ*_SRreceived_(*t*+1) represents the TD-RPE received by (both subsystems of) the SR-based system (see below), and also a constant decay/forgetting (with the rate *d* = 0.001) at every time-step:

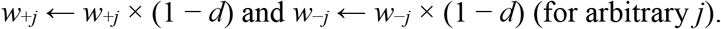

The SR-based system specific value *V*_SR_(*S*(*t*)) was calculated as:

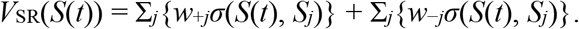

The mean (average) of the IR and SR-system-specific values was calculated as the integrated value *V*(*S*), which was used for action selection in the same manner as in the original model. Notably, because the amount of value update was proportional to the multiplication of the learning rate (*a*_SR+_ or *a*_SR−_) and the SR feature variable *σ*(*S*(*t*), *S*_*j*_), which was presumably represented in the cortex, the learning rate here can represent, instead of the literal learning rate, the strength of cortical input to the striatal direct or indirect pathway (or in other words, the connection strength between the cortical neurons/regions hosting SR and the particular striatal pathway). Similarly, the learning rate *a*_IR+_ or *a*_IR−_ can represent the connection strength between the cortical neurons/regions hosting IR and the striatal direct or indirect pathway, respectively.

When the agent was at state *S*(*t*) at time-step *t* and moved to *S*(*t*+1) at *t*+1, IR- and SR-value-specific TD-RPEs (*δ*_IR_(*t*+1) and *δ*_SR_(*t*+1)) were calculated based on the IR- and SR-system-specific values, respectively:

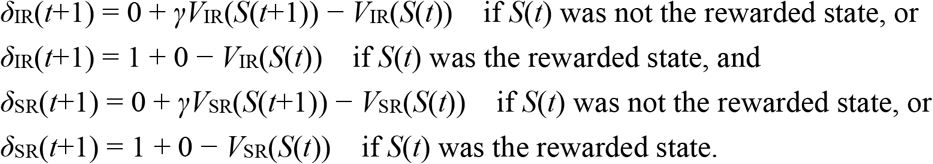

Then, TD-RPEs *received* by the IR- and SR-based systems (*δ*_IRreceived_(*t*+1) and *δ*_SRreceived_(*t*+1)) were calculated as follows:

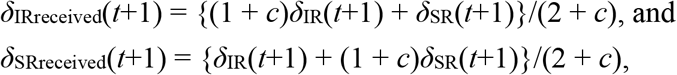

where *c* indicates the bias that the IR-based system receives more IR-value-specific TD-RPEs than SR-value-specific TD-RPEs and vice versa, and was varied to 0, 0.1, 0.2, or 0.3.

### Simulations and statistics

Simulations were conducted *n* = 100 times for the reward navigation tasks, or 1000 times for the two-stage tasks, for each condition by using MATLAB. 50 sets of 100 simulations were conducted for Figure 3C (including the one set shown in Figure 3A,B) and Figure S3D. Probabilities and pseudorandom numbers were implemented by “rand”, “randn”, and “randperm” functions of MATLAB. Standard deviation (SD) was calculated with normalization by *n*, and standard error of the mean (SEM) was approximately calculated as SD/√*n*.

### Code availability

Codes to generate the data presented in the figures and tables are available at: https://github.com/kenjimoritagithub/sr3

## Supplementary Text

### More comprehensive examination of the parameter space

In the simulations of the original reward navigation task by the original two-systems models in the main text, we examined the cases where the sum of the learning rates from positive and negative TD-RPEs (*α*_+_ + *α*_−_) in both systems was one of the three values (0.75, 1, or 1.25) and the learning rate for the update of SR features was fixed at 0.05. In order to examine the parameter space more widely, we varied the learning rate from positive or negative TD-RPEs in each of the two systems freely from 0.2, 0.35, 0.5, 0.65, or 0.8 and the learning rate for the update of SR features from 0.05, 0.1, 0.15, 0.2, or 0.25 (only for the cases where at least one of the two systems employed SR), also varying the time discount factor (*γ* = 0.6, 0.7, or 0.8) and the degree of choice exploration-exploitation (i.e., inverse temperature) (*β* = 5, 10, or 15), for each of the cases where the two systems employed SR and IR, SR only, or IR only. Figure S1 shows the mean performance of the model consisting of SR- and IR-based systems. Each row of panels in Figure S1 shows the mean performance for each set of time discount factor and inverse temperature (shown in the left), varying the learning rate for the update of SR features (shown in the top), projected onto the plane consisting of the ratios of the learning rates from positive and negative TD-RPEs in the two systems (*α*_SR+_/*α*_SR−_ and *α*_IR+_/*α*_IR−_); there were 25 cases with *α*_SR+_/*α*_SR−_ = *α*_IR+_/*α*_IR−_ = 1, which were difficult to draw on the single point (1, 1) in this projected plane and thus omitted. Figure S2 shows the mean performance of the model consisting of two SR-based systems or two IR-based systems, projected onto the plane consisting of *α*_1+_/*α*_1−_ and *α*_2+_/*α*_2−_; 25 cases with *α*_1+_/*α*_1−_ = *α*_2+_/*α*_2−_ = 1 were omitted for the same reason as above. Table S1 shows the sets of learning rate parameters that gave top ten mean performance for each set of time discount factor and inverse temperature (shown in the left) in the model consisting of SR- and IR-based systems (left), two SR-based systems (middle), and two IR-based systems (right). For the model consisting of SR- and IR-based systems, combinations of appetitive SR-based system (*α*_+_/*α*_−_ > 1) and aversive IR-based system (*α*_+_/*α*_−_ < 1) are shown in bold italic.

Regarding the model consisting of SR- and IR-based systems (Figure S1 and Table S1), under the examined conditions differing in the degrees of temporal discounting and/or choice exploration-exploitation (each row of panels in Figure S1 and sub-tables in Table S1), combination of appetitive SR-based and aversive IR-based systems (shown in bold italic in Table S1) with relatively small learning rate for SR feature update (*α*_SRfeature_) generally achieved good performance. As *α*_SRfeature_ increased, combination of the *α*_+_/*α*_−_ ratios in the two systems that achieved good performance approached to the vertical diagonal line (i.e., similar *α*_+_/*α*_−_ ratios in both systems), but the achieved good performance itself tended to decrease when *α*_SRfeature_ exceeded 0.1 or 0.15.

**Figure S1.**
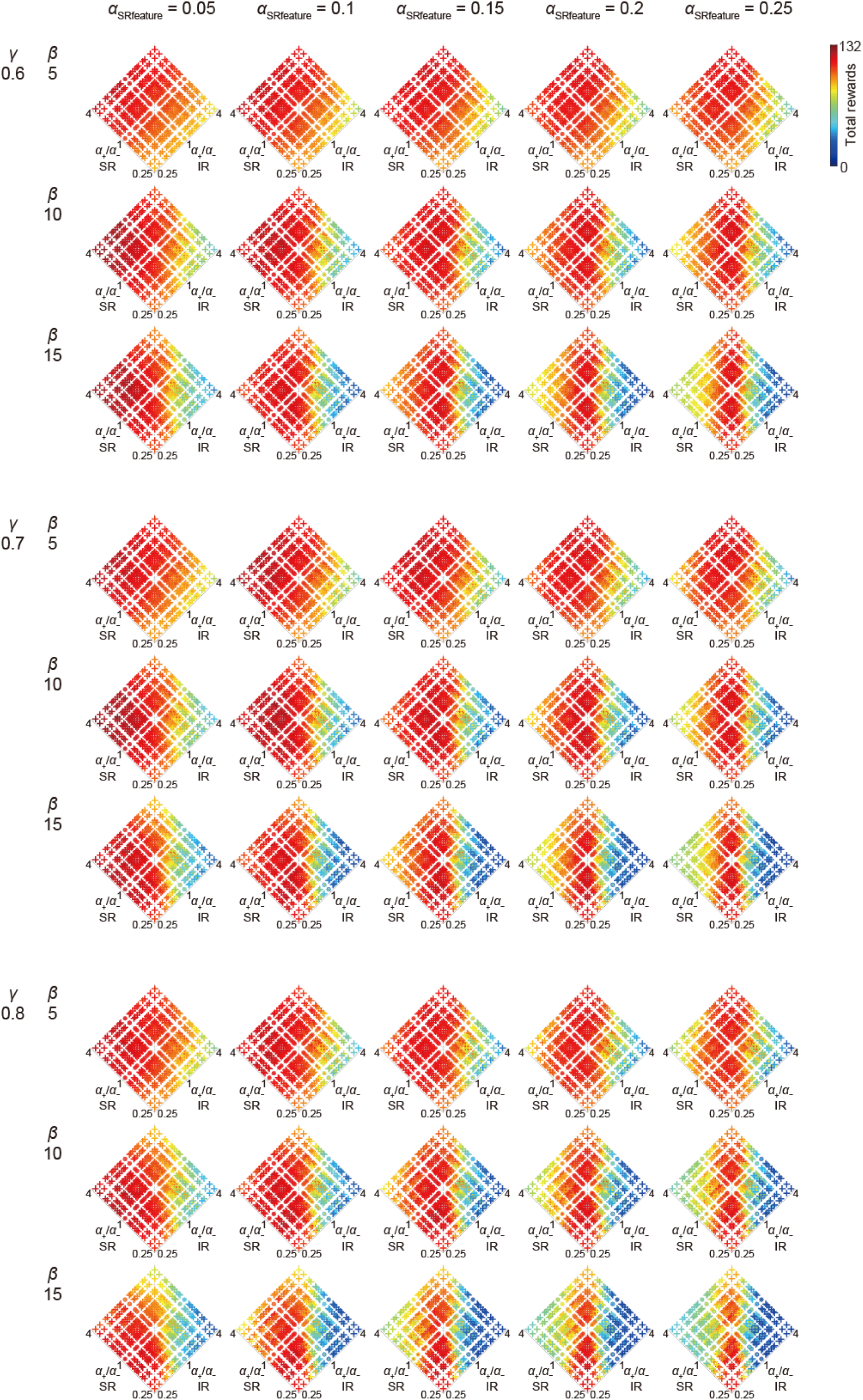
Performance of the model consisting of SR-based and IR-based systems in a broader parameter space. Each row of panels shows the mean performance for each set of time discount factor (*γ*) and inverse temperature (*β*) (shown in the left), varying the learning rate for the update of SR features (*α*_SRfeature_, shown in the top), projected onto the plane consisting of the ratios of the learning rates from positive and negative TD-RPEs in the two systems (*α*_SR+_/*α*_SR−_ and *α*_IR+_/*α*_IR−_). There were 25 cases with *α*_SR+_/*α*_SR−_ = *α*_IR+_/*α*_IR−_ = 1, which were difficult to draw on the single point (1, 1) in this projected plane and thus omitted. In the cases with *α*_SR+_/*α*_SR−_ = 1 or *α*_IR+_/*α*_IR−_ = 1 but not *α*_SR+_/*α*_SR−_ = *α*_IR+_/*α*_IR−_ = 1, there were five cases having the same set of (*α*_SR+_/*α*_SR−_, *α*_IR+_/*α*_IR−_) (because *α*_+_/*α*_−_ = 1 corresponds to five cases with *α*_+_ = *α*_−_ = 0.2, 0.35, 0.5, 0.65, or 0.8), and these five cases were drawn by concentric circles with larger radius corresponding to larger learning rate. In the cases other than the special cases mentioned just above, each case was drawn by a cross. The color of the symbols (cross or circle) indicates the mean performance, in reference to the color bar in the top right.

**Table S1.**
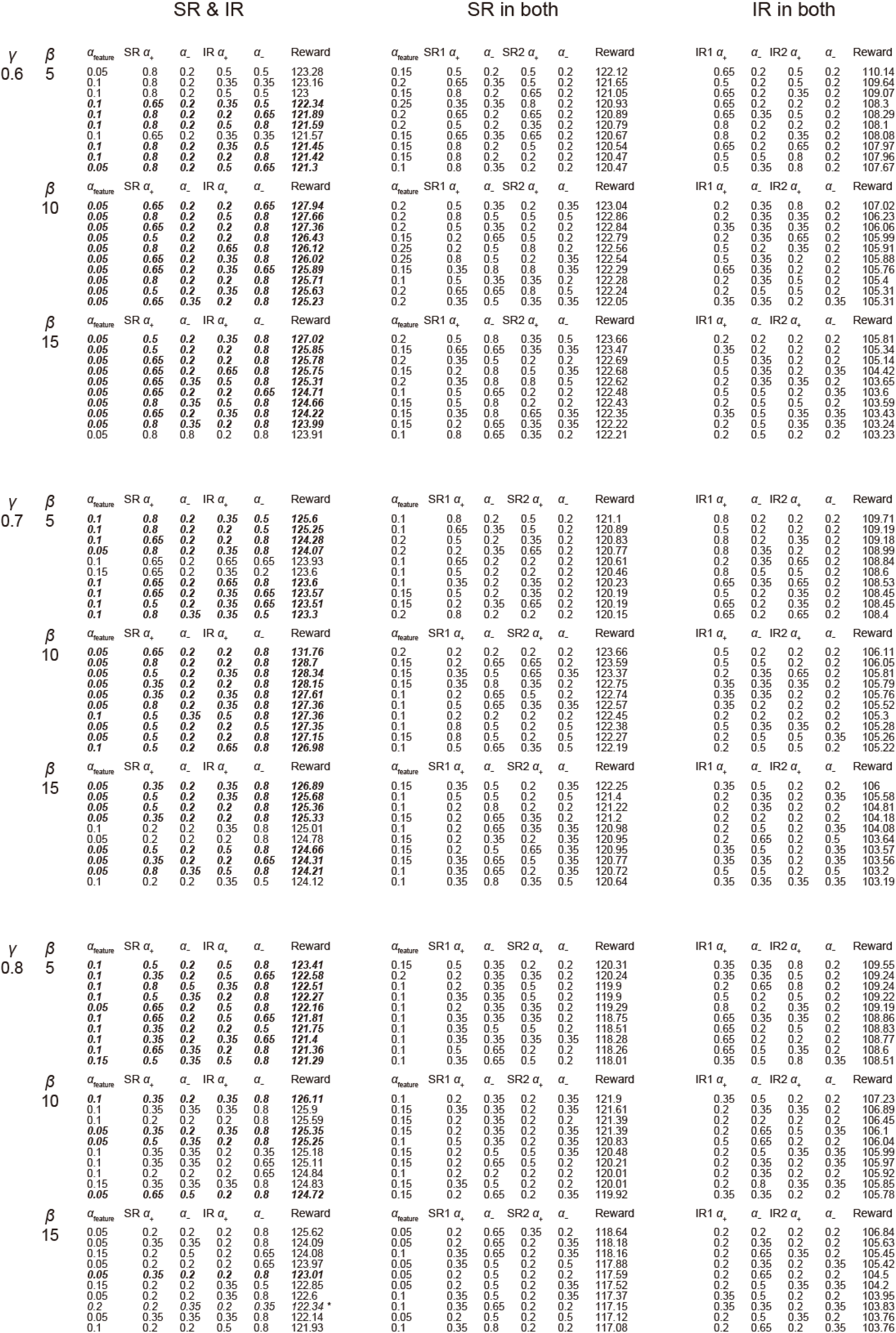
Performance of the models consisting of two systems in a broader parameter space. The left, middle, and right part shows the results for the model consisting of SR- and IR-based systems, two SR-based systems, and two IR-based systems, respectively. Each sub-table shows the sets of learning rate parameters that gave top ten mean performance for each set of time discount factor (*γ*) and inverse temperature (*β*) (shown in the left) in each of the three models. For the model consisting of SR- and IR-based systems, cases with a combination of appetitive (*α*_+_/*α*_−_ > 1) SR-based system and aversive (*α*_+_/*α*_−_ < 1) IR-based system are shown in bold italic; notably, even in all the other cases shown in this table except for a case shown in italic with asterisk in the right, the *α*_+_/*α*_−_ ratio was higher in the SR-based system than in the IR-based system.

Regarding the models where both systems employed only SR or IR (Figure S2 and Table S1), the SR-only models generally outperformed the IR-only models. For the SR-only model, increase in *α*_SRfeature_ up to 0.15 or 0.2 improved the performance in many cases, but further increase appears to be not generally beneficial. As for the SR-only model, the case with (*γ, β*) = (0.6, 15), performed well, comparably to the case with (*γ, β*) = (0.7, 10). So we further examined cases with (*γ, β*) = (0.5, 15), (0.5, 20), and (0.6, 20). Table S2 shows the sets of learning rates giving top ten mean performance for these three cases, which look largely comparable to the cases (*γ, β*) = (0.6, 15) or (0.7, 10). As for the IR-only model, the cases with small inverse temperature (*β* = 5) performed well. So we further examined cases with *β* = 2.5, but performance was generally worse than the cases with *β* = 5 (best mean performance was 107.2, 107.76, and 105.12 in the cases with *γ* = 0.6, 0.7, and 0.8, respectively).

**Figure S2.**
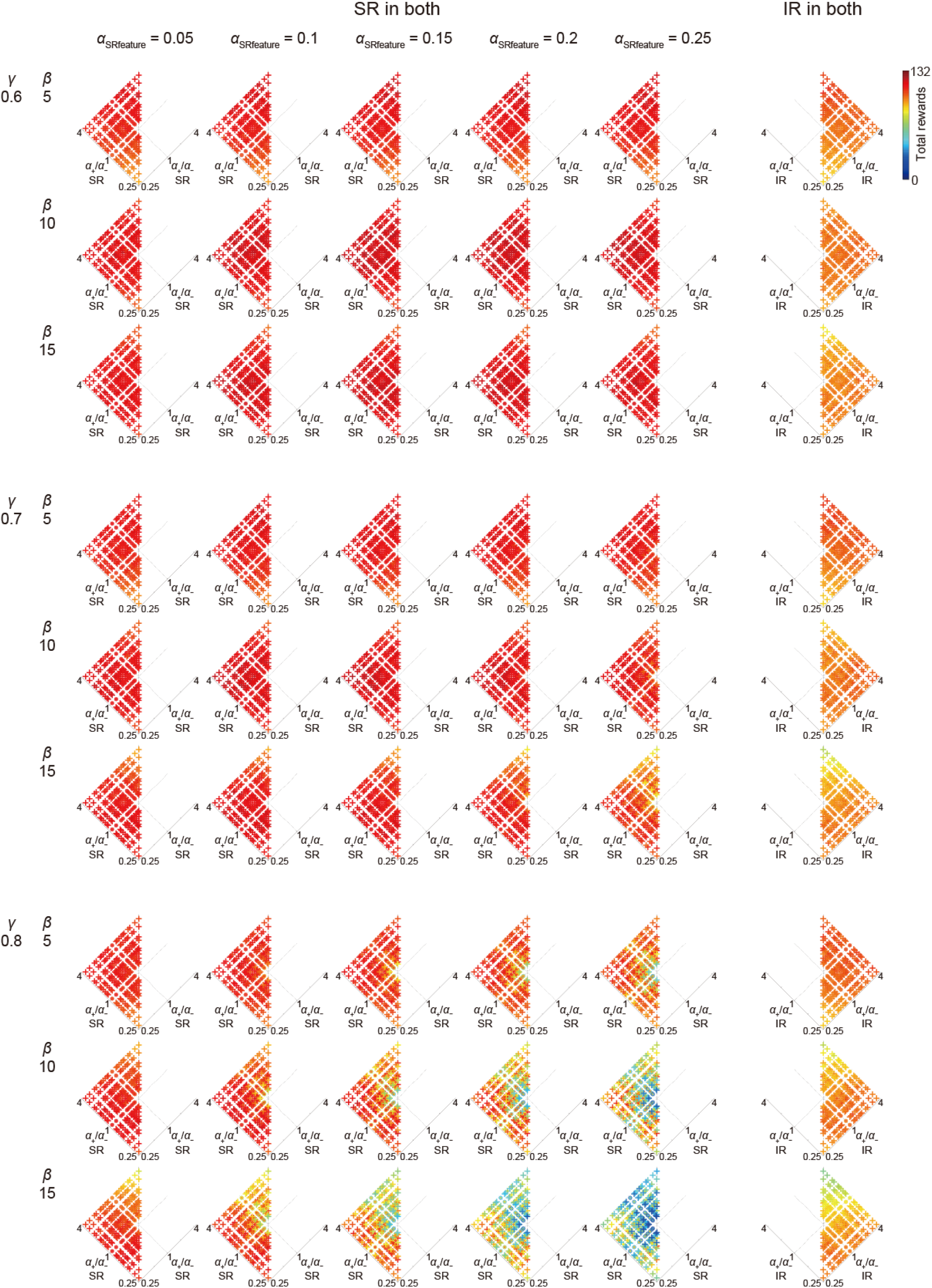
Performance of the model consisting of two SR-based systems or two IR-based systems in a broader parameter space. Each row of panels shows the mean performance for each set of time discount factor (*γ*) and inverse temperature (*β*) (shown in the left), varying the learning rate for the update of SR features (*α*_SRfeature_, shown in the top) for the cases with two SR-based systems (five panels from the left), projected onto the plane consisting of the ratios of the learning rates from positive and negative TD-RPEs in the two systems (*α*_1+_/*α*_1−_ and *α*_2+_/*α*_2−_). The cases with *α*_1+_/*α*_1−_ = *α*_2+_/*α*_2−_ = 1 were not drawn, and the cases with *α*_1+_/*α*_1−_ = 1 or *α*_2+_/*α*_2−_ = 1 but not *α*_1+_/*α*_1−_ = *α*_2+_/*α*_2−_ = 1 were drawn by concentric circles, and all the other cases were drawn by crosses, in a similar manner to Figure S1. The color of the symbols (cross or circle) indicates the mean performance, in reference to the color bar in the top right. Only the left or right side is shown for the cases with two SR- or IR-based systems, respectively, for the same reason as in Figure 6.

**Table S2.** Results of additional simulations for the model consisting of two SR-based systems. Each sub-table shows the sets of learning rate parameters that gave top ten mean performance for each set of time discount factor (*γ*) and inverse temperature (*β*) shown in the left.

| $\gamma$ | $\beta$ | $\alpha_{\text{feature}}$ | SR1 $\alpha_+$ | $\alpha_-$ | SR2 $\alpha_+$ | $\alpha_-$ | Reward |
| --- | --- | --- | --- | --- | --- | --- | --- |
| 0.5 | 15 | 0.2 | 0.35 | 0.35 | 0.35 | 0.35 | 123.65 |
|  |  | 0.1 | 0.8 | 0.8 | 0.35 | 0.35 | 123.04 |
|  |  | 0.15 | 0.35 | 0.65 | 0.5 | 0.5 | 122.88 |
|  |  | 0.25 | 0.5 | 0.65 | 0.8 | 0.5 | 122.88 |
|  |  | 0.15 | 0.5 | 0.5 | 0.35 | 0.2 | 122.79 |
|  |  | 0.1 | 0.65 | 0.8 | 0.35 | 0.2 | 122.65 |
|  |  | 0.2 | 0.65 | 0.65 | 0.8 | 0.35 | 122.48 |
|  |  | 0.15 | 0.35 | 0.5 | 0.35 | 0.35 | 122.28 |
|  |  | 0.25 | 0.5 | 0.8 | 0.8 | 0.5 | 122.25 |
|  |  | 0.2 | 0.65 | 0.8 | 0.5 | 0.2 | 122.17 |
| $\gamma$ | $\beta$ | $\alpha_{\text{feature}}$ | SR $\alpha_+$ | $\alpha_-$ | IR $\alpha_+$ | $\alpha_-$ | Reward |
| 0.5 | 20 | 0.2 | 0.2 | 0.8 | 0.8 | 0.2 | 123.58 |
|  |  | 0.25 | 0.5 | 0.65 | 0.65 | 0.2 | 123.53 |
|  |  | 0.2 | 0.65 | 0.5 | 0.2 | 0.5 | 123.09 |
|  |  | 0.2 | 0.8 | 0.65 | 0.2 | 0.2 | 122.99 |
|  |  | 0.15 | 0.5 | 0.8 | 0.65 | 0.5 | 122.33 |
|  |  | 0.2 | 0.65 | 0.65 | 0.2 | 0.2 | 122.28 |
|  |  | 0.2 | 0.2 | 0.65 | 0.65 | 0.35 | 122.26 |
|  |  | 0.2 | 0.5 | 0.8 | 0.5 | 0.5 | 122.2 |
|  |  | 0.25 | 0.2 | 0.65 | 0.35 | 0.35 | 122.03 |
|  |  | 0.25 | 0.8 | 0.8 | 0.2 | 0.5 | 122.02 |
| $\gamma$ | $\beta$ | $\alpha_{\text{feature}}$ | SR $\alpha_+$ | $\alpha_-$ | IR $\alpha_+$ | $\alpha_-$ | Reward |
| 0.6 | 20 | 0.15 | 0.65 | 0.5 | 0.2 | 0.5 | 124 |
|  |  | 0.1 | 0.2 | 0.5 | 0.5 | 0.35 | 121.91 |
|  |  | 0.15 | 0.35 | 0.8 | 0.2 | 0.2 | 121.71 |
|  |  | 0.2 | 0.2 | 0.65 | 0.8 | 0.5 | 121.66 |
|  |  | 0.15 | 0.2 | 0.5 | 0.2 | 0.2 | 121.59 |
|  |  | 0.05 | 0.2 | 0.65 | 0.35 | 0.5 | 121.46 |
|  |  | 0.1 | 0.2 | 0.35 | 0.2 | 0.35 | 121.4 |
|  |  | 0.1 | 0.35 | 0.8 | 0.2 | 0.35 | 121.33 |
|  |  | 0.1 | 0.2 | 0.65 | 0.8 | 0.2 | 121.31 |
|  |  | 0.2 | 0.2 | 0.35 | 0.2 | 0.35 | 121.31 |

Comparing the three types of models using SR and IR, SR only, or IR only, under each of the nine (3×3) sets of time discount factor and inverse temperature (*γ* = 0.6, 0.7, 0.8 and *β* = 5, 10, 15; shown in Figures S1 and S2 and Table S1) that were examined for all the three types of models, best mean performance, among the examined sets of learning rate parameters, was achieved by a combination of SR- and IR-based systems. Under seven out of the nine sets, combination of appetitive SR- and aversive IR-based systems achieved the best performance, and even under the remaining two, best performance was achieved by a combination where the ratio of learning rates from positive over negative TD-RPEs was higher in the SR-based system than in the IR-based system.

### Immediate sensitivity to outcome devaluation of the appetitive SR and aversive IR combination

A popular experimental paradigm to test whether animal’s behavior is goal-directed/model-based or not is the outcome devaluation (Balleine and O’Doherty, 2010; Dolan and Dayan, 2013). Animals first learn that a particular action leads to a particular reward. Then, the reward is specifically devalued, for example, by making the animals satiated with food reward. Thereafter, the animals’ tendency to take that action is examined. Immediate decrease in the tendency, compared with the case without devaluation, indicates that the animals estimated the action’s value taking the devaluation of its outcome into account, without experiencing the association of the action and the devalued reward, i.e., behaved in a goal-directed/model-based manner. Agent adopting SR-based TD learning can also exhibit immediate sensitivity to outcome devaluation even though it does not have explicit model of the environment, because the environmental structure is partially embedded in the SR. It should depend on negative TD-RPEs generated when the agent experienced the devaluation of the reward. Then, an emerging question is whether agent having the combination of appetitive SR- and aversive IR-based systems, which performed well in our tasks, can also exhibit devaluation sensitivity despite the limited SR-based aversive learning.

In order to examine this, we simulated a devaluation test (Figure S3A,B). There were two goal states, Goal 1 at (3, 1) and Goal 2 at (1, 3), and when the agent reached one of the pre-goal states, i.e., (2, 1) or (1, 2), in the next time step it always proceeded to the corresponding goal state unless it was at the end of epoch. There was an initial 500 time-steps no reward epoch, followed by a 500 time-steps learning epoch, where two rewards were introduced at the two goal states. During the learning epoch, when the agent reached either goal state, it obtained the reward, and reward was immediately resupplied. At the beginning of the learning epoch, there was a forced learning period, in which the agent forcedly experienced each reward (transition from pre-goal state to goal state) 10 times, and after that, the agent freely behaved according to learned state values. Agent then experienced devaluation of one of the rewards, specifically, the reward introduced at Goal 2 (1, 3). The agent was directly placed at this goal state, experiencing TD error-based updates but without experiencing state transition. After this devaluation phase, agent was placed at (1, 1), and which of the two pre-goal states the agent initially reached was examined.

**Figure S3.**
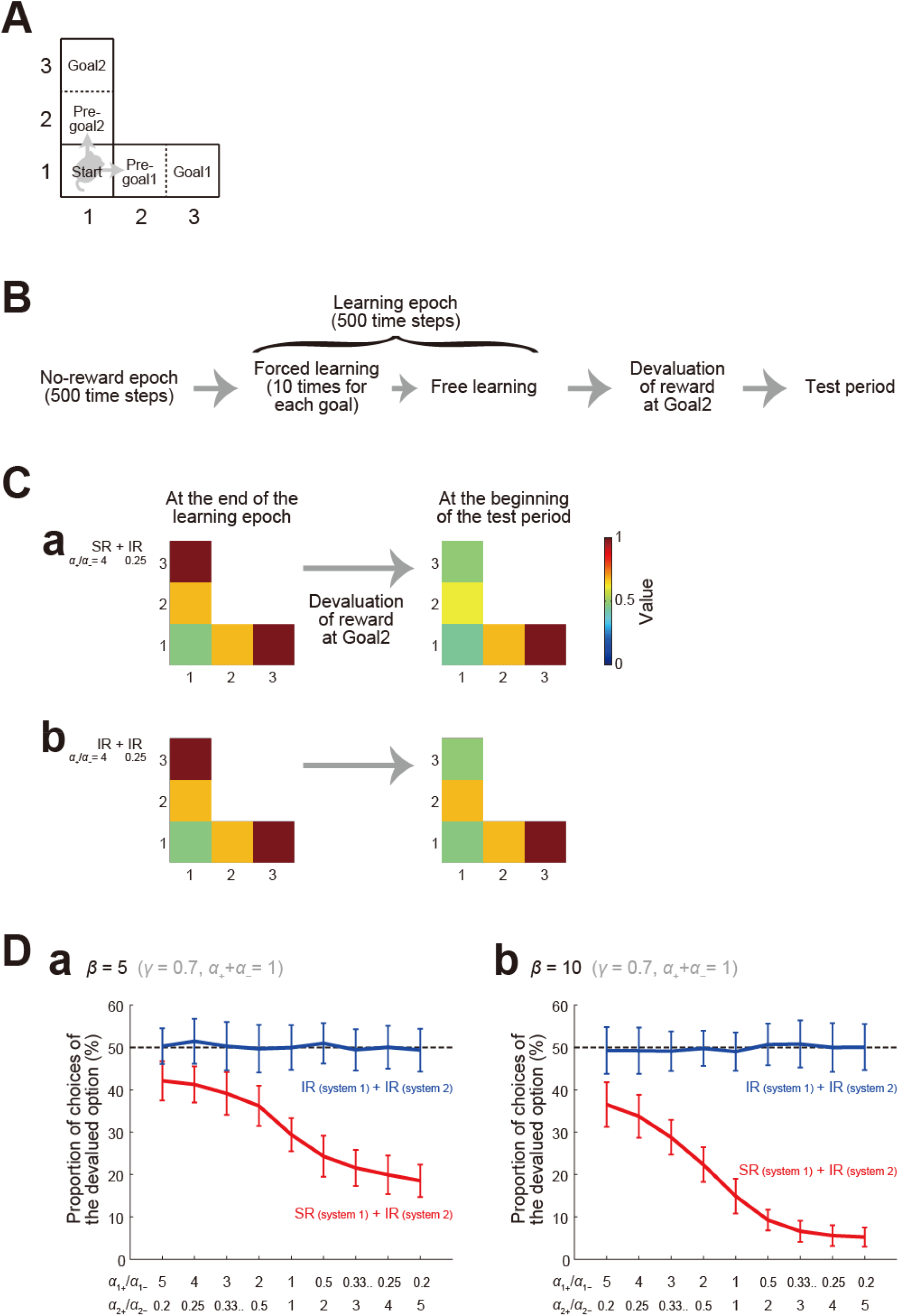
Immediate sensitivity to outcome devaluation of the agent equipped with two systems. **(A)** A narrowed grid space for the simulated outcome devaluation test. There were two goal states, Goal 1 at (3, 1) and Goal 2 at (1, 3), and when the agent reached one of the pre-goal states, i.e., (2, 1) or (1, 2), in the next time step it always proceeded to the corresponding goal state unless it was at the end of task epoch. **(B)** Temporal structure of the simulated outcome devaluation test. There was an initial 500 time-steps no-reward epoch, followed by a 500 time-steps learning epoch. At the beginning of the learning epoch, there was a forced learning period, in which the agent forcedly experienced each reward (transition from pre-goal state to goal state) 10 times. The learning epoch was followed by a devaluation phase, in which the agent experienced devaluation of the reward at Goal 2 (1, 3). Then, in the test period, the agent was placed at (1, 1), and which of the two pre-goal states the agent initially reached was examined. **(C) (a)** Examples of the integrated state values at the end of the learning epoch (left panels) and at the beginning of the test period following devaluation (right panels) for agent with appetitive SR-based system (*α*_+_/*α*_−_ = 5) and aversive IR-based system (*α*_+_/*α*_−_ = 1/5) **(b)** Examples for agent with appetitive and aversive IR-based systems (*α*_+_/*α*_−_ = 5 and 1/5). **(D) (a)** The mean proportion of reaching the pre-goal state preceding the devalued Goal 2 in the test period in the cases with SR- and IR-based systems (red lines) and with two IR-based systems (blue lines) with different *α*_+_/*α*_−_ ratios (horizontal axis) and *β* = 5. For each condition, proportion in 100 simulations was calculated for 50 sets, and its mean and standard deviation were plotted. **(b)** Cases with *β* = 10.

We examined the behavior of agent equipped with combination of SR-based and IR-based systems, with the ratios of the learning rates from positive and negative TD-RPEs in the two systems varied while their multiplication was kept constant at 1 (i.e., conditions corresponding to those on the horizontal diagonal in Figure 3A). As a control, we also examined the behavior of agent equipped with two IR-based systems, with the ratios of the learning rates from positive and negative TD-RPEs in the two systems similarly varied. The sum of positive- and negative TD-RPE-based learning rates, the degree of exploitation/exploration, and the time discount factor were set as same as those for Figure 3 (*α*_+_ + *α*_−_ = 1, *β* = 5, and *γ* = 0.7); *β* = 10 was also examined.

Figure S3C shows examples of the integrated state values at the end of the learning epoch and at the beginning of the test period following devaluation for agent with appetitive SR-based system (*α*_+_/*α*_−_ = 5) and aversive IR-based system (*α*_+_/*α*_−_ = 1/5) (Figure S3Ca) or with appetitive and aversive IR-based systems (*α*_+_/*α*_−_ = 5 and 1/5) (Figure S3Cb). In both cases, at the end of the learning epoch (left panels), the values of the two pre-goal states leading to the two goal states were both learned to become high. In the case of agent with appetitive SR- and aversive IR-based systems, devaluation of reward at Goal 2 caused a decrease in the value of Goal 2 itself and also a decrease in the value of the pre-goal state leading to Goal 2 (Figure S3Ca, right panel). In contrast, in the case of agent with appetitive and aversive IR-based systems, reward devaluation only decreased the value of Goal 2 itself and did not change the value of the pre-goal state (Figure S3Cb, right panel).

Figure S3D shows the mean proportion of reaching the pre-goal state preceding the devalued Goal 2 in the test period in the cases with SR- and IR-based systems (red lines) and with two IR-based systems (blue lines) with different *α*_+_/*α*_−_ ratios (horizontal axis); cases with *β* = 5 and *β* = 10 are shown in Figures S3Da and S3Db, respectively. For each condition, proportion in 100 simulations was calculated for 50 sets, and its mean and standard deviation were plotted. In the cases with two IR-based systems, the mean proportion was nearly 50% in all the cases, indicating that the agent reached either of the pre-goal states regardless of whether the upcoming reward was devalued or not, i.e., failed to exhibit immediate sensitivity to outcome devaluation. By contrast, in the cases with SR- and IR-based systems, the mean proportion was lower than 50% in all the cases, although the degree of deviation from 50% decreased as the *α*_+_/*α*_−_ ratio for the SR-based system increased. Also, deviation was greater for the case with *β* = 10 than for the case with *β* = 5. These results indicate that even the agent with appetitive SR- and aversive IR-based system can exhibit immediate sensitivity to outcome devaluation at least to a certain extent despite the limited SR-based learning from negative TD-RPEs.

## Acknowledgements

KM was supported by Grant-in-Aid for Scientific Research (No. 20H05049) of the Ministry of Education, Culture, Sports, Science and Technology in Japan (MEXT) (http://www.mext.go.jp/en/). YK was supported by Grant-in-Aid for Scientific Research (No. 17H06311, 20H03359) of the MEXT and the Japan Society for the Promotion of Science (https://www.jsps.go.jp/english/) and by the Cooperative Study Program of National Institute for Physiological Sciences (https://www.nips.ac.jp/eng/) (Section of Electron Microscopy).

## References

Balleine, B.W., and O’Doherty, J.P. (2010). Human and rodent homologies in action control: corticostriatal determinants of goal-directed and habitual action. Neuropsychopharmacology 35, 48–69. 10.1038/npp.2009.131.

Ballion, B., Mallet, N., Bézard, E., Lanciego, J.L., and Gonon, F. (2008). Intratelencephalic corticostriatal neurons equally excite striatonigral and striatopallidal neurons and their discharge activity is selectively reduced in experimental parkinsonism. Eur J Neurosci 27, 2313-2321. EJN6192 [pii] 10.1111/j.1460-9568.2008.06192.x.

Barreto, A., Dabney, W., Munos, R., Hunt, J.J., Schaul, T., van Hasselt, H., and Silver, D. (2016). Successor Features for Transfer in Reinforcement Learning. 1606.05312.

Bogacz, R. (2020). Dopamine role in learning and action inference. Elife 9. 10.7554/eLife.53262.

Chen, Y., Monaco, S., Byrne, P., Yan, X., Henriques, D.Y., and Crawford, J.D. (2014). Allocentric versus egocentric representation of remembered reach targets in human cortex. J Neurosci 34, 12515–12526. 10.1523/JNEUROSCI.1445-14.2014.

Collins, A.G., and Frank, M.J. (2014). Opponent actor learning (OpAL): modeling interactive effects of striatal dopamine on reinforcement learning and choice incentive. Psychol Rev 121, 337–366. 10.1037/a0037015.

Cui, G., Jun, S.B., Jin, X., Pham, M.D., Vogel, S.S., Lovinger, D.M., and Costa, R.M. (2013). Concurrent activation of striatal direct and indirect pathways during action initiation. Nature 494, 238–242. 10.1038/nature11846.

Daw, N.D., Gershman, S.J., Seymour, B., Dayan, P., and Dolan, R.J. (2011). Model-based influences on humans’ choices and striatal prediction errors. Neuron 69, 1204-1215. S0896-6273(11)00125-5 [pii] 10.1016/j.neuron.2011.02.027.

Dayan, P. (1993). Improving Generalization for Temporal Difference Learning: The Successor Representation. Neural Computation 5, 613–624.

Dolan, R.J., and Dayan, P. (2013). Goals and habits in the brain. Neuron 80, 312–325. 10.1016/j.neuron.2013.09.007.

Frank, M.J., Seeberger, L.C., and O’reilly, R.C. (2004). By carrot or by stick: cognitive reinforcement learning in parkinsonism. Science 306, 1940–1943.1102941 [pii] 10.1126/science.1102941.

Gardner, M.P.H., Schoenbaum, G., and Gershman, S.J. (2018). Rethinking dopamine as generalized prediction error. Proc Biol Sci 285, 20181645. 10.1098/rspb.2018.1645.

Garvert, M.M., Dolan, R.J., and Behrens, T.E. (2017). A map of abstract relational knowledge in the human hippocampal-entorhinal cortex. Elife 6, e17086. 10.7554/eLife.17086.

Gershman, S.J., Moore, C.D., Todd, M.T., Norman, K.A., and Sederberg, P.B. (2012). The successor representation and temporal context. Neural Comput 24, 1553–1568. 10.1162/NECO_a_00282.

Groman, S.M., Keistler, C., Keip, A.J., Hammarlund, E., DiLeone, R.J., Pittenger, C., Lee, D., and Taylor, J.R. (2019a). Orbitofrontal Circuits Control Multiple Reinforcement-Learning Processes. Neuron 103, 734-746.e733. 10.1016/j.neuron.2019.05.042.

Groman, S.M., Massi, B., Mathias, S.R., Curry, D.W., Lee, D., and Taylor, J.R. (2019b). Neurochemical and Behavioral Dissections of Decision-Making in a Rodent Multistage Task. J Neurosci 39, 295–306. 10.1523/JNEUROSCI.2219-18.2018.

Hamid, A.A., Frank, M.J., and Moore, C.I. (2021). Wave-like dopamine dynamics as a mechanism for spatiotemporal credit assignment. Cell 184, 2733-2749.e2716. 10.1016/j.cell.2021.03.046.

Hart, G., Bradfield, L.A., and Balleine, B.W. (2018a). Prefrontal Corticostriatal Disconnection Blocks the Acquisition of Goal-Directed Action. J Neurosci 38, 1311–1322. 10.1523/JNEUROSCI.2850-17.2017.

Hart, G., Bradfield, L.A., Fok, S.Y., Chieng, B., and Balleine, B.W. (2018b). The Bilateral Prefronto-striatal Pathway Is Necessary for Learning New Goal-Directed Actions. Curr Biol 28, 2218-2229.e2217. 10.1016/j.cub.2018.05.028.

Hikida, T., Kimura, K., Wada, N., Funabiki, K., and Nakanishi, S. (2010). Distinct roles of synaptic transmission in direct and indirect striatal pathways to reward and aversive behavior. Neuron 66, 896-907. S0896-6273(10)00379-X [pii] 10.1016/j.neuron.2010.05.011.

Igarashi, K.M., Ieki, N., An, M., Yamaguchi, Y., Nagayama, S., Kobayakawa, K., Kobayakawa, R., Tanifuji, M., Sakano, H., Chen, W.R., and Mori, K. (2012). Parallel mitral and tufted cell pathways route distinct odor information to different targets in the olfactory cortex. J Neurosci 32, 7970–7985. 10.1523/JNEUROSCI.0154-12.2012.

Iino, Y., Sawada, T., Yamaguchi, K., Tajiri, M., Ishii, S., Kasai, H., and Yagishita, S. (2020). Dopamine D2 receptors in discrimination learning and spine enlargement. Nature 579, 555–560. 10.1038/s41586-020-2115-1.

Kim, H.F., Amita, H., and Hikosaka, O. (2017). Indirect Pathway of Caudal Basal Ganglia for Rejection of Valueless Visual Objects. Neuron 94, 920-930.e923. 10.1016/j.neuron.2017.04.033.

Kool, W., Cushman, F.A., and Gershman, S.J. (2016). When Does Model-Based Control Pay Off? PLoS Comput Biol 12, e1005090. 10.1371/journal.pcbi.1005090.

Kravitz, A.V., Tye, L.D., and Kreitzer, A.C. (2012). Distinct roles for direct and indirect pathway striatal neurons in reinforcement. Nat Neurosci 15, 816–818. nn.3100 [pii] 10.1038/nn.3100.

Kress, G.J., Yamawaki, N., Wokosin, D.L., Wickersham, I.R., Shepherd, G.M., and Surmeier, D.J. (2013). Convergent cortical innervation of striatal projection neurons. Nat Neurosci 16, 665–667. 10.1038/nn.3397.

Lee, R.S., Engelhard, B., Witten, I.B., and Daw, N.D. (2022). A vector reward prediction error model explains dopaminergic heterogeneity. bioRxiv https://doi.org/10.1101/2022.02.28.482379.https://doi.org/10.1101/2022.02.28.482379.

Lee, S.J., Lodder, B., Chen, Y., Patriarchi, T., Tian, L., and Sabatini, B.L. (2021). Cell-type-specific asynchronous modulation of PKA by dopamine in learning. Nature 590, 451–456. 10.1038/s41586-020-03050-5.

Lehnert, L., Tellex, S., and Littman, M.L. (2017). Advantages and Limitations of using Successor Features for Transfer in Reinforcement Learning. arXiv, 1708.00102v00101.

Lei, W., Jiao, Y., Del Mar, N., and Reiner, A. (2004). Evidence for differential cortical input to direct pathway versus indirect pathway striatal projection neurons in rats. J Neurosci 24, 8289–8299. 24/38/8289 [pii] 10.1523/JNEUROSCI.1990-04.2004.

Lu, J., Cheng, Y., Xie, X., Woodson, K., Bonifacio, J., Disney, E., Barbee, B., Wang, X., Zaidi, M., and Wang, J. (2021). Whole-Brain Mapping of Direct Inputs to Dopamine D1 and D2 Receptor-Expressing Medium Spiny Neurons in the Posterior Dorsomedial Striatum. eNeuro 8. 10.1523/ENEURO.0348-20.2020.

Martiros, N., Kim, S.E., Kapoor, V., and Murthy, V.N. (2021). Distinct representation of cue-outcome association by D1 and D2 neurons in the olfactory striatum. bioRxiv, https://doi.org/10.1101/2021.1111.1101.466363.https://doi.org/10.1101/2021.11.01.466363.

Matamales, M., McGovern, A.E., Mi, J.D., Mazzone, S.B., Balleine, B.W., and Bertran-Gonzalez, J. (2020). Local D2-to D1-neuron transmodulation updates goal-directed learning in the striatum. Science 367, 549–555. 10.1126/science.aaz5751.

Mikhael, J.G., and Bogacz, R. (2016). Learning Reward Uncertainty in the Basal Ganglia. PLoS Comput Biol 12, e1005062. 10.1371/journal.pcbi.1005062.

Mikhael, J.G., Kim, H.R., Uchida, N., and Gershman, S.J. (2022). The role of state uncertainty in the dynamics of dopamine. Curr Biol 32, 1077-1087.e1079. 10.1016/j.cub.2022.01.025.

Momennejad, I., Russek, E.M., Cheong, J.H., Botvinick, M.M., Daw, N.D., and Gershman, S.J. (2017). The successor representation in human reinforcement learning. Nat Hum Behav 1, 680–692. 10.1038/s41562-017-0180-8.

Morishima, M., Morita, K., Kubota, Y., and Kawaguchi, Y. (2011). Highly differentiated projection-specific cortical subnetworks. J Neurosci 31, 10380–10391.

Morita, K. (2014). Differential cortical activation of the striatal direct and indirect pathway cells: reconciling the anatomical and optogenetic results by using a computational method. J Neurophysiol 112, 120–146. 10.1152/jn.00625.2013.

Morita, K., Im, S., and Kawaguchi, Y. (2019). Differential Striatal Axonal Arborizations of the Intratelencephalic and Pyramidal-Tract Neurons: Analysis of the Data in the MouseLight Database. Front Neural Circuits 13, 71. 10.3389/fncir.2019.00071.

Morita, K., and Kato, A. (2014). Striatal dopamine ramping may indicate flexible reinforcement learning with forgetting in the cortico-basal ganglia circuits. Front Neural Circuits 8, 36. 10.3389/fncir.2014.00036.

Morita, K., and Kawaguchi, Y. (2019). A Dual Role Hypothesis of the Cortico-Basal-Ganglia Pathways: Opponency and Temporal Difference Through Dopamine and Adenosine. Front Neural Circuits 12, 111. 10.3389/fncir.2018.00111.

Möller, M., and Bogacz, R. (2019). Learning the payoffs and costs of actions. PLoS Comput Biol 15, e1006285. 10.1371/journal.pcbi.1006285.

Nonomura, S., Nishizawa, K., Sakai, Y., Kawaguchi, Y., Kato, S., Uchigashima, M., Watanabe, M., Yamanaka, K., Enomoto, K., Chiken, S., et al. (2018). Monitoring and Updating of Action Selection for Goal-Directed Behavior through the Striatal Direct and Indirect Pathways. Neuron. https://doi.org/10.1016/j.neuron.2018.08.002.

Peak, J., Chieng, B., Hart, G., and Balleine, B.W. (2020). Striatal direct and indirect pathway neurons differentially control the encoding and updating of goal-directed learning. Elife 9. 10.7554/eLife.58544.

Piray, P., and Daw, N.D. (2021). Linear reinforcement learning in planning, grid fields, and cognitive control. Nat Commun 12, 4942. 10.1038/s41467-021-25123-3.

Reiner, A., Hart, N.M., Lei, W., and Deng, Y. (2010). Corticostriatal projection neurons -dichotomous types and dichotomous functions. Front Neuroanat 4, 142. 10.3389/fnana.2010.00142.

Russek, E.M., Momennejad, I., Botvinick, M.M., Gershman, S.J., and Daw, N.D. (2017). Predictive representations can link model-based reinforcement learning to model-free mechanisms. PLoS Comput Biol 13, e1005768. 10.1371/journal.pcbi.1005768.

Russek, E.M., Momennejad, I., Botvinick, M.M., Gershman, S.J., and Daw, N.D. (2021). Neural evidence for the successor representation in choice evaluation. bioRxiv https://doi.org/10.1101/2021.08.29.458114.https://doi.org/10.1101/2021.08.29.458114.

Shen, W., Flajolet, M., Greengard, P., and Surmeier, D.J. (2008). Dichotomous dopaminergic control of striatal synaptic plasticity. Science 321, 848-851. 321/5890/848 [pii] 10.1126/science.1160575.

Stachenfeld, K.L., Botvinick, M.M., and Gershman, S.J. (2017). The hippocampus as a predictive map. Nat Neurosci 20, 1643–1653. 10.1038/nn.4650.

Sutton, R., and Barto, A. (1998). Reinforcement Learning (MIT Press).

Tai, L.H., Lee, A.M., Benavidez, N., Bonci, A., and Wilbrecht, L. (2012). Transient stimulation of distinct subpopulations of striatal neurons mimics changes in action value. Nat Neurosci 15, 1281–1289. 10.1038/nn.3188.

Town, S.M., Brimijoin, W.O., and Bizley, J.K. (2017). Egocentric and allocentric representations in auditory cortex. PLoS Biol 15, e2001878. 10.1371/journal.pbio.2001878.

Wall, N.R., De La Parra, M., Callaway, E.M., and Kreitzer, A.C. (2013). Differential innervation of direct-and indirect-pathway striatal projection neurons. Neuron 79, 347–360. 10.1016/j.neuron.2013.05.014.

Wang, C., Chen, X., and Knierim, J.J. (2020). Egocentric and allocentric representations of space in the rodent brain. Curr Opin Neurobiol 60, 12–20. 10.1016/j.conb.2019.11.005.

Watabe-Uchida, M., Eshel, N., and Uchida, N. (2017). Neural Circuitry of Reward Prediction Error. Annu Rev Neurosci 40, 373–394. 10.1146/annurev-neuro-072116-031109.

Winnubst, J., Bas, E., Ferreira, T.A., Wu, Z., Economo, M.N., Edson, P., Arthur, B.J., Bruns, C., Rokicki, K., Schauder, D., et al. (2019). Reconstruction of 1,000 Projection Neurons Reveals New Cell Types and Organization of Long-Range Connectivity in the Mouse Brain. Cell 179, 268-281.e213. 10.1016/j.cell.2019.07.042. (End)

